# Chemical Editing of Proteoglycan Architecture

**DOI:** 10.1101/2021.04.02.437933

**Authors:** Timothy O’Leary, Meg Critcher, Tesia N. Stephenson, Xueyi Yang, Noah H. Bartfield, Richard Hawkins, Mia L. Huang

**Author notes:** Equal authorship (Timothy O’Leary, Meg Critcher).

## Abstract

Proteoglycans are heterogeneous macromolecular glycoconjugates that orchestrate many important cellular processes. While much attention has focused on the poly-sulfated glycosaminoglycan chains that decorate proteoglycans, other important elements of proteoglycan architecture, such as their core proteins and cell surface localization, have garnered less emphasis. Hence, comprehensive structure-function relationships that consider the replete proteoglycan architecture as glycoconjugates are limited. Here, we present a comprehensive approach to study proteoglycan structure and biology by fabricating defined semi-synthetic modular proteoglycans that can be tailored for cell surface display. To do so, we integrate amber codon reassignment in the expression of sequence-fined proteoglycan core proteins, metabolic oligosaccharide engineering to produce functionalizable glycosaminoglycans, and bioorthogonal click chemistry to covalently tether the two components. These materials permit the methodical dissection of the parameters required for optimal binding and function of various proteoglycan-binding proteins, and they can be modularly displayed on the surface of any living cell. We demonstrate that these sophisticated materials can recapitulate the functions of native proteoglycans in mouse embryonic stem cell differentiation and cancer cell spreading, while permitting the identification of the most important contributing elements of proteoglycan architecture toward function. This technology platform will confer structural resolution toward the investigation of proteoglycan structure-function relationships in cell biology.

## Introduction

Proteoglycans (PGs) are ubiquitous macromolecular glycoconjugates that serve dynamic and important roles in cell biology (1). There is incredible detail embedded within the PG architecture, as it is composed of a core protein covalently decorated with poly-sulfated glycosaminoglycans (GAGs). Moreover, the conjugate product can be differentially displayed as soluble or cell surface-anchored molecules (**Fig. 1A**). Despite the overwhelming presence of PGs at the glycocalyx (2), the systematic exploration of the roles of PG architecture as replete glycoconjugates is lacking. Because of the polymorphic nature of PGs, current approaches focus on achieving structural resolution of the individual components (either the protein or GAG chains), but in doing so, they sacrifice the total architecture of PGs.

**Figure 1.**
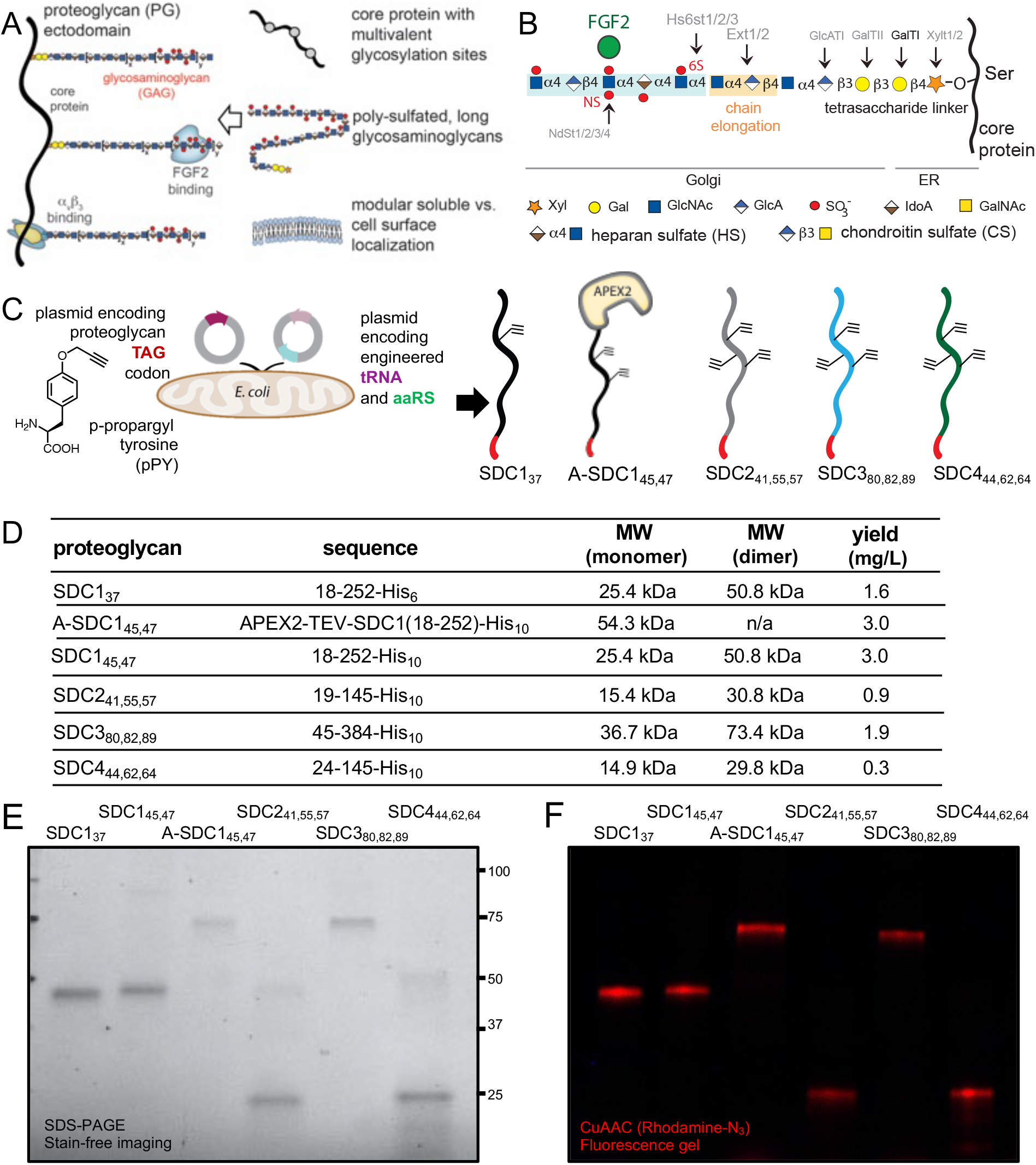
General proteoglycan architecture and biosynthesis. (**A**) Proteoglycan architecture is composed of multiple elements, a core protein is covalently decorated with glycosaminoglycan (GAG) chains, and the entire glycoconjugate can be differentially displayed as soluble or cell surface-anchored molecules. (**B**) The ribosomally-synthesized core protein is enzymatically modified with a tetrasaccharide (GlcA-Gal-Gal-Xyl) linker and further elongated (by Exostosin 1/2, Ext1/2). Changes in disaccharide composition and linkages result in the various classes of GAGs (e.g. heparan, chondroitin, dermatan, keratan sulfate). Sulfotransferases (e.g. Ndsts, Hs6sts) transfer sulfate groups to GAGs, shown here are sulfotransferases specific for heparan sulfate PGs. GAG chains are heterogeneous in size and composition, and they can encode specificity for growth factors, such as Fibroblast Growth Factor-2 (FGF2). (**C**) Amber codon reassignment is used to produce p-propargyltyrosine (pPY)-modified SDC1-4 core proteins. SDC1 cDNA containing modifications at GAG attachment sites to the TAG amber codon were co-transformed in *E. coli* along with a plasmid that encodes for a re-engineered tRNA/aaRS pair, resulting in cells expressing SDC ectodomains with pPY at 1-3 sites (positions denoted by subscripts). (**D**) Table summarizing the PG ectodomain sequences, expected molecular weights, and expression yields. (**E**) SDS-PAGE analysis of the expressed PGs. (**F**) Copper-catalyzed azide-alkyne cycloadditions (CuAAC) with rhodamine-azide shows the presence of fluorescent protein bands, indicative of successful pPY incorporation.

PG biogenesis begins with the ribosomal synthesis of the core protein (**Fig. 1B**), which is composed of an extracellular-facing ectodomain as well as transmembrane and cytoplasmic domains. The core protein is then post-translationally modified with a tetrasaccharide linker (GlcA-Gal-Gal-Xyl) at serine/threonine residues and extensively elongated by the action of glycosyltransferases, which sequentially install monosaccharides to the growing chain. The elongated chain is composed of repeating units of an amino sugar (GlcNAc or GalNAc) and an uronic acid (glucuronic or iduronic acid), where the disaccharide compositions and glycosidic linkages dictate whether the GAG is classified as a heparan sulfate (HS), chondroitin sulfate (CS), dermatan sulfate (DS), or keratan sulfate (KS). The GAG chains are further modified with sulfate groups at various positions (e.g., 2-, 3-, and 6-*O*, or *N*-sulfation) by sulfotransferases. In total, an elaborate symphony of >40 enzymes are responsible for generating polydisperse GAGs (3).

PG glycoconjugates transact with their environment via their extracellular-facing ectodomains, which carry out a majority of their functions. Although the covalently attached GAGs can dominate PG sizes, the core protein itself deserves attention. Core proteins can complex with their own set of unique interactors (e.g. receptor tyrosine kinases to activate signaling cascades) that are distinct from its family members, and some ligands even prefer to bind deglycosylated PGs (4). Recent work has also demonstrated that the core protein can even dictate the extent of GAG sulfation (5). From a structural perspective, the core proteins also organize the GAG chains into multivalent copies, and this arrangement can be important for function. The decorating GAGs also serve important roles, where different sulfation patterns can encode binding motifs for GAG-binding proteins, including Fibroblast Growth Factor-2 (FGF2) to regulate cell growth and differentiation. (6, 7) Through variation of these elements, PGs can shape a range of functions and cell behavior. For example, a PG called syndecan-1 (SDC1) is present at similar levels on both simple and stratified epithelial cells, yet because they exhibit various GAG glycosylation states, they can act as rheostats to meet the vastly different demands of the epithelium to achieve diversified cell adhesion. (8)

Efficient methods to access defined and complete PGs that permit a holistic way to modulate their architectures, are currently lacking. Although advances in the chemical and chemoenzymatic synthesis of carbohydrates have provided well-defined GAG structures (9), these techniques often require substantial technical expertise in carbohydrate chemistry, and to date, only short GAG oligomers (~10 residues) (10, 11) or glycopeptides (12) have been made. Such short oligomers fail to span the chemical space occupied by large GAG chains (100-500 monosaccharide residues). While heparin is commonly used as a surrogate molecule for HS, their structures differ quite dramatically. Native HS is composed of domains of sulfated and non-sulfated regions and is much longer and less sulfated compared to heparin (~0.6 vs. 2.6 sulfates/disaccharide). Moreover, while heparin is often used in its soluble format, native HS is present as both a soluble or a cell surface-anchored molecule, and this differing presentation can have profound impact on function. Neither the oligomers, glycopeptides, nor heparin recapitulate the compartmentalized domain arrangement of native GAGs, where regions of sulfation flank the non-sulfated regions. Failure to control the sulfation patterns of GAGs can lead to contradictory experimental observations. (13, 14) Macromolecular synthetic polymers that can present glycans in a multivalent manner to span larger distances have been employed to recreate GAG functions. (15–17) However, these approaches overlook the functional roles of the core protein. (15, 18) Additionally, simplified genetic techniques to manipulate PG architecture may fail to provide a complete picture of PG biology. Targeting the expression of PG core proteins may result in dysregulated GAG chains, whereas targeting the GAG biosynthetic machinery can alter PG expression patterns or even lethality (19). Moreover, manipulation of sheddases, the enzymes that cleave cell surface-anchored ectodomains to soluble fragments, often results in the off-target removal of many other proteins (20).

Here, we present a straightforward platform to control all of the components of a PG ectodomain, the core protein, the appended GAGs, and its membrane localization (**Fig. 1A**). This platform, rooted in chemical biology, offers unique opportunities to study the biological functions of PGs in high molecular detail, while maintaining a focus on the structures of the individual components. To generate functionalizable PG ectodomains, we chose to use unnatural amino acid (UAA) incorporation to install site-specific bioorthogonal reactive handles. In order to produce materials that recapitulate the native sizes and domain arrangement of native GAGs, we hijacked the cellular biosynthetic machinery using synthetic azidoxylosides. Finally, to tune the display of the edited PGs onto cell surfaces, we employed a metal-based cell surface engineering strategy. Our engineered PGs have permitted a renewed understanding of how the presence of GAGs or the core proteins influence the binding of cytokines and extracellular matrix components, as well as showcase the importance of properly considering the membrane localization of PGs. We used the engineered SDC1 ectodomains to systematically study interactions with the α_v_β_3_ integrin or FGF2 and catalogued their corresponding effects towards mouse embryonic stem cell (mESCs) differentiation and cancer cell spreading.

## Results

### Production of functionalizable ectodomains through protein engineering

We have developed a modular platform that permits the tailored installation of GAGs and membrane anchors onto PG core proteins. Our platform (**Fig. 1C**) entails the recombinant expression of poly-histidine-tagged core protein ectodomains in *E. coli* hosts, where serine/threonine GAG glycosylation sites have been modified to alkyne-containing unnatural amino acids, such as *p*-propargyl tyrosine (pPY). This modification is accomplished by re-assigning the amber codon, TAG, to encode for pPY through engineered tRNA and corresponding aminoacyl tRNA synthetases (aaRS) (21). We co-transformed competent Rosetta DE3 *E. coli* with individual plasmids encoding mouse syndecans 1-4 (SDC1-4) ectodomains and the suppressor plasmid, pULTRA-CNF (Addgene # 48215) (21). This plasmid encodes for an orthogonal tRNA/aaRS pair derived from *Methanococcus jannaschii*. As a proof of concept, we expressed pPY-containing SDC core protein ectodomains with C-terminal poly-histidine (poly-His) tags for nickel-nitriloacetic acid (NTA)-based purification (22) or complexation (23). We prepared pPY-modified versions of all four members of the syndecan family (SDC1-4), with each protein incorporating one to three pPY amino acids at residues known to incorporate either a HS or CS GAG (**Fig. 1C, Fig. 1D**; pPY positions depicted by subscripts). The SDCs are a family of type I transmembrane core proteins (5), where each SDC bears unique ectodomain sequences, of which SDC1 is the most studied within the family (24). SDC1 is comprised of three conserved sites for GAG attachment; position 37 is known to be modified by CS (sometimes HS), whereas residues 45 and 47 are thought to be mostly modified by HS, and its multivalent arrangement can be required for function. (25) We prepared a monovalent version of SDC1 with one pPY site at residue 37, SDC1_37_ (**Fig. S1**), and we also explored the possibility of creating fusion proteins of SDC1 for various applications. Hence, we also prepared a APEX2 peroxidase-conjugated version of SDC1 bearing two pPY sites at residues 45 and 47, separated by a TEV protease-cleavable tag (A-SDC1_45,47_; **Fig. S2A, B**). The divalent SDC1_45,47_ was prepared by treatment of the APEX2 conjugate with TEV protease (**Fig. S2C**). We focused our efforts towards the three conserved GAG-modified residues within the SDC family, as most functions are ascribed to them (25). As expected, all SDC ectodomains were expressed in their de-glycosylated states (*E. coli* does not include the pertinent GAG biosynthesis machinery; **Fig. S3**). (20) These proteins were efficiently purified from lysates using Ni/NTA-based chromatography, and except for A-SDC1_45,47_, they migrated on a SDS-PAGE gel as dimers, consistent with previous reports (**Fig. 1E**). (20) We confirmed the presence of the pPY amino acids by copper(I)-catalyzed [3+2] cycloaddition (CuAAC) (26) with a TAMRA azide fluorophore (**Fig. 1F**). The pPY-modified SDCs were expressed in moderate to high yields (0.3-3.0 mg/L), even with three proximally-located pPY sites. We also confirmed that SDC1_37_ retained its capacity to be selectively processed by known sheddases, as it was processed by MMP-9 and not by ADAMTS-1 (**Fig. S4**). Using the same pULTRA-CNF plasmid, we also expressed modest yields of another UAA-modified proteoglycan, glypican-1 (GPC1), with three of its known HS-modified sites (residues 485, 487, 489) mutated to another propargyl-lysine (**Fig. S5**).

### Metabolic engineering with azido xylosides to hijack GAG production

To generate GAGs that recapitulate the length and domain organization of native biomolecules, we prepared recombinant HS (rHS) and CS (rCS) GAGs using a metabolic oligosaccharide engineering strategy, where we hijacked GAG biosynthesis (**Fig. 1B**) using synthetic small molecules (27). Previous work has shown that xyloside molecules derivatized with hydrophobic aglycones, such as phenyl groups, can efficiently serve as decoys and compete for native GAG elongation processes (28–31). We posited that this strategy could be co-opted towards the production of GAGs with functionalizable handles at the reducing ends, by simply appending the hydrophobic moieties with azide groups. Starting from a bromo-xyloside intermediate and *p*-azidophenol, we generated *p-*azidophenol-xyloside **1**(**Fig. 2A; Fig S6**). We modeled how compound **1** docks into the β4GalT7 galactosyltransferase active sites and observed that it minimizes to −6.3 kcal/mol (**Fig. 2B**), similar to the phenyl derivative lacking the azide group (**Fig. S7**). Furthermore, **1** is positioned such that it can be accessed for subsequent elongation in the GAG biosynthesis pathway. These experiments gave us confidence that the azide group minimally perturbs the aglycone, and that compound **1** could potentially be used to produce recombinant GAGs from cells.

**Figure 2.**
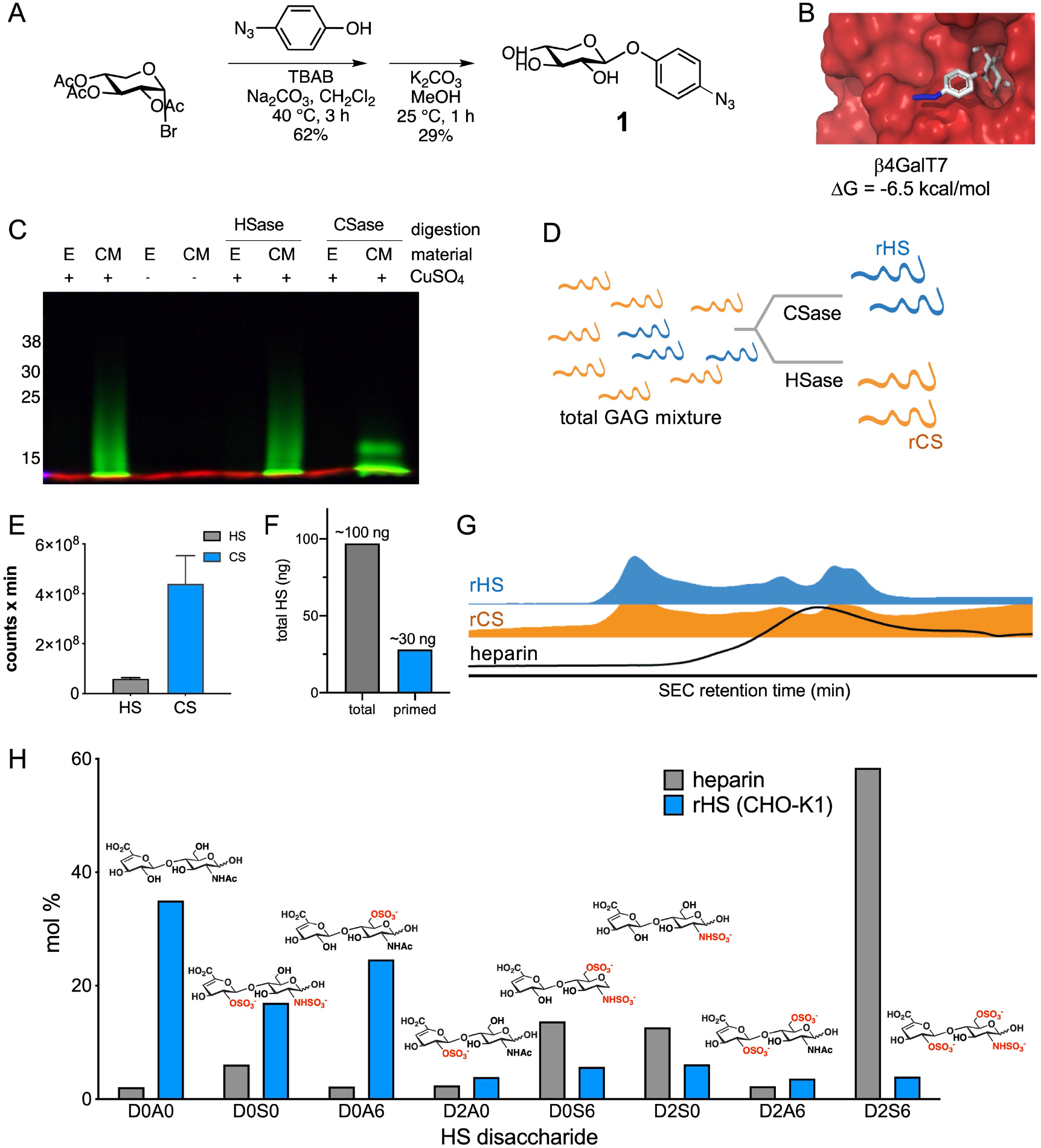
Azidoxyloside 1 hijacks GAG biosynthesis to produce recombinant GAGs. (**A**) Synthetic strategy to produce azidoxyloside **1.** (**B**) Docking of the enzyme acceptor β4GalT7 with compound **1**. (**C**) By incubating mammalian cells with azidoxyloside **1** (400 μM), we generated recombinant GAGs that bear functional azide handles at the reducing ends. Fluorescence SDS-PAGE gel confirms that GAGs isolated from conditioned media (CM) retain the azide functional handle (confirmed via CuAAC with AF488-alkyne). Most of the GAGs were recovered from the conditioned media (CM) versus cell extracts (E). (**D**) Selective enzymatic digestion of the total GAG mixture with chondroitinase (CSase) or heparinases (HSase) was used to obtain either recombinant HS (rHS) or recombinant CS (rCS), respectively. (**E**) As expected from CHO-K1 cells, the GAG mixture is mostly composed of CS. Bar graphs depict means and standard deviations. (**F**) About 30% of total HS isolated from CHO-K1 cells bear the azide functional group. (**G**) Recombinant azide-primed HS differs in charge and size compared to heparin. (**H**) Recombinant HS bears GAG composition akin to native HS and significantly differs from heparin. Here, are shown the top eight most common motifs found in HS.

Thus, we incubated **1** (400 μM) with wild-type Chinese hamster ovary (CHO)-K1 cells (**Fig. 2C**), a cell line commonly used for GAG priming studies, and observed minimal effects in cell viability. We observed significant enhancements in azide-primed GAGs with **1** when incubation times were increased from 48 to 72 hours (**Fig. S8A**). After incubation, both the conditioned medium (CM) and cell extracts (E) were collected. To verify the presence of the azide group, the isolated materials were reacted with AlexaFluor488-alkyne using CuAAC and the resulting products analyzed by SDS-PAGE (**Fig. 2C**). Upon fluorescence visualization, we observed the presence of broad and diffuse fluorescent smears migrating from 8-30 kDa, indicating extensive molecular heterogeneity. As expected, the fluorescent GAGs were mostly present in the CM, as expected, with little to no fluorescent materials was observed in cell extracts (**Fig. S8B**). This observation is consistent with the reactions where copper was excluded, confirming that the fluorescence signal arose due to CuAAC.

When we selectively digested the total recombinant GAGs (rGAGs) with either a cocktail of heparinases (HSase) or chondroitinases (CSase; **Fig. 2D**), we observed that CSase digestion yielded visibly smaller molecular weight fluorescent fragments, while HSase digestion did not cause an immediately obvious change in the migration of the fluorescent band (**Fig. 2C**). This SDS-PAGE experiment suggested that majority of the material from CHO cells was composed of CS, consistent with the *Ext1*-hemizygous nature of CHO cells. Moreover, it suggests that the recombinant GAGs generated from azidoxyloside **1** faithfully captures host cell GAG production. Indeed, when we quantified the composition of the material from the CM, we observed that >80% is composed of CS (**Fig. 2E**) and that ~30% of total HS included the azide motif (**Fig. 2F**). The high concentrations of compounds required for priming are consistent with the proposed metabolic engineering strategy. Although only ~30% of the total HS GAGs are equipped with the azide motif, the unprimed non-azide tagged GAGs are inert towards subsequent CuAAC, and the isolated CM can be used without further purification.

We compared the sizes of the rGAGs isolated from CHO-K1 cells to heparin and observed that the recombinant materials differed in size (**Fig. 2G**). While rHS and rCS seemed to migrate similarly by size exclusion chromatography, both have shorter retention times compared to heparin, suggesting their comparatively large sizes (**Fig. 2G**). Although strategies to sequence HS GAGs remain limited, their disaccharide compositions can be used as a metric to fingerprint their identities (**Fig. S9**). Thus, we also analyzed the composition of the rHS from CHO-K1 by enzymatic depolymerization into the repeating disaccharide units (**Fig. 2H**) (32) As anticipated, the non-sulfated D0A0 disaccharide motif accounted for a large part of these rHS GAGs (33). This resulting composition is expected of native HS and is consistent with its domain-like arrangement, where regions of non-sulfated D0A0 disaccharides flank the sulfated regions. This disaccharide sulfation pattern is in stark contrast to that of heparin, which is mostly composed of the tri-sulfated D2S6 disaccharide motif. We also observed similar HS and CS disaccharide compositions in the rHS and rCS compared to native HS and CS produced by the CHO-K1 cells (**Fig. S10**), as well as in rCS from 48 and 72 hours of priming (**Fig. S11**). Overall, these observations indicate that the priming process produces GAG of similar composition to native host cell GAGs.

### CuAAC generates proteoglycan glycoconjugates

To construct GAG-decorated PG ectodomains, we subjected azide-terminating GAGs, azido-heparin or azido-CS (**Fig. S13**) or our recombinant GAGs (**Fig. 2**) to CuAAC with the pPY-modified SDC1 ectodomain SDC1_37_. After one hour, we analyzed the crude reactions using anion exchange chromatography and observed significant conversion to the higher molecular weight and more anionic GAG-conjugated product (**Fig. 3A**). We consistently observed high conversion yields (~90%) regardless of the GAG, core protein, or number of conjugation sites (**Fig. S14-S16**). The reaction conversions and migration patterns of the conjugates were similar to those of a model pPY-modified green fluorescent protein (GFP; **Fig. S17**). We analyzed the fractions from the reaction product of SDC1_37_ conjugated to total rGAGs from CHO-K1 (generated via 48 or 72 hr incubation periods) and observed SDC1-positive bands using an anti-SDC1 antibody (clone 281-2), which displays reactivity towards both the GAG conjugate and to a lesser extent, the core protein (**Fig. 3B**). In contrast to the localized band for the SDC1_37_ core protein, the glycoconjugates appeared as large, diffuse signals spanning molecular weights up to 260 kDa when probed using 281-2 or anti-HS antibody clone 10E4 (**Fig. 3C**). Through selective digestion of the rGAG mixture (**Fig. 2D**) and subsequent CuAAC conjugation to SDC1_37_ or SDC1_45,47_, we also prepared SDC1_37rHS_ and SDC1_45,47rHS_. Performing the conjugation step in vitro and purification of the resulting conjugates by anion exchange and size exclusion chromatography further ensures that the purified materials are free of copper to mitigate potential toxicity concerns in live cells. Overall, we demonstrate that despite the poly-anionic nature of the GAGs and the SDC1 ectodomain (pI = 4.7), CuAAC is a robust method to generate PG glycoconjugates.

**Figure 3.**
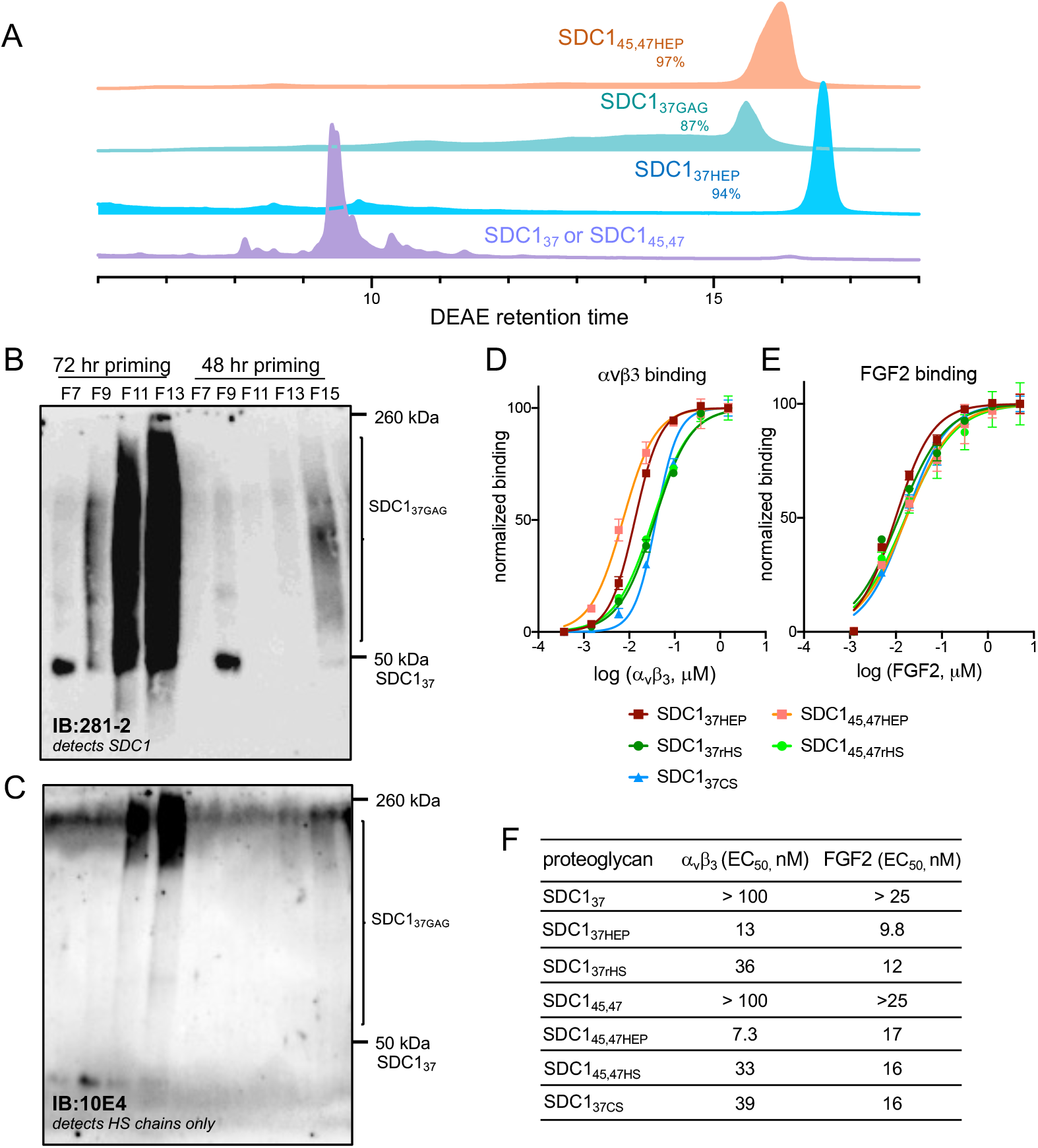
CuAAC generates replete proteoglycan glycoconjugates. (**A**) pPY-modified ectodomains can be efficiently reacted in high conversion yields with GAGs (e.g. azide-heparin, rGAGs from CHO-K1 cells) using CuAAC. Similar traces and yields were obtained with rHS and rCS obtained from cells. (**B**) Fractions collected from purification of the glycoconjugates resulting from recombinant GAGs primed for 48 vs 72 hours were further analyzed by Western blotting. Probing for SDC1 using an anti-SDC1 antibody (clone 281-2) demonstrates greatly enhanced priming with longer incubation. Although we expect a product of ~85 kDa (25 kDa core protein + 60 kDa (recombinant GAG), the large smears (50-260 kDa) are likely due to the unusual migration of these conjugates in SDS-PAGE. Clone 281-2 displays weak affinity for the core protein, which is enhanced upon conjugation with any GAG. (**C**) Probing the same blot for the presence of HS-modified conjugates using an anti-HS antibody (clone 10E4) demonstrates that despite its low abundance in the mixture relative to CS, it can still be conjugated to SDC1_37_. (**D**) ELISA with immobilized SDC1 ectodomains against recombinant α_v_β_3_ or (**E**) FGF2. (**F**) Tabulated apparent affinity constants (EC_50_) shown for each PG and target.

### PG editing enables systematic dissection of structure-binding relationships

We first analyzed whether our PGs had discernable differences in binding to known SDC1-binding proteins, such as the α_v_β_3_ integrin (34) or FGF2 (35). Using an ELISA with the engineered SDC1 ectodomains immobilized onto the plate, we observed that α_v_β_3_ integrin (**Fig. 3D; Table S2**) and FGF2 (**Fig. 3E; Table S3**) display overall greater affinity for GAG-conjugated SDC1_37_ compared to the non-conjugated ectodomain alone. Effective binding constants (EC_50_; **Fig. 3F**) measured for the conjugates against α_v_β_3_ followed the ranking: SDC1_45,47HEP_ < SDC1_37HEP_ < SDC1_45,47HS_ < SDC1_37HS_ < SDC1_37CS_. The α_v_β_3_ integrin has been predicted to bind the SDC1 core protein via residues 88-121 (36) and more recently, to heparin and HS via a series of basic lysine residues from both the α_v_ and the β_3_ chains (37). The most potent binder in the series, the divalent SDC1_45,47HEP_ conjugate displayed an approximately 2-fold lower apparent affinity constant relative to the monovalent SDC1_37HEP_ (EC_50_: 7.3 vs. 13 nM). This observation suggests that the extent of binding is highly influenced by the number of GAG chains, as similarly observed in other studies (25). The higher affinities observed for the heparin conjugates compared to rHS suggests that the extent of sulfation, or the presence of 3-*O*-sulfation in heparin, also strongly enhances affinity. The GAG conjugates displayed significantly greater affinity for FGF2 as well, similar to α_v_β_3_. EC_50_ values derived for binding FGF2 ranked in the order SDC1_37HEP_ < SDC1_37rHS_ < SDC1_45,47HS_ ~ SDC1_37CS_ < SDC1_45,47HEP_. Notably, we observed an increase in affinity for the divalent SDC1_45,47HEP_ relative to its monovalent counterpart SDC1_37HEP_ (EC_50_: 9.8 vs. 17 nM).

### Cell surface engineering and stem cell differentiation

We next explored whether altered cell surface presentations (*i.e.* soluble vs. membrane-anchored) altered SDC1-mediated effects in a relevant biological system. To create cell surface-anchored SDC1 ectodomains, we employed a cell surface engineering approach (**Fig. 4A**), where live cells are first incubated with a lipid functionalized with NTA (e.g. cholesterol-PEG2000-NTA; cholPEGNTA) before incubation with SDC1 PGs in the presence of Ni(OAc)_2_ (38). The affinity of the C-terminal poly-His tag for the tetradentate Ni-NTA chelating ligand results in complexation and the presentation of the SDC1 ectodomain at the cell surface. We used cholesterol as a lipid anchor, because of its high solubility, its ability to passively incorporate into any phospholipid membrane and to recycle back to the cell surface upon endocytosis (39). We chose to use mouse embryonic stem cells (mESCs) as the backdrop for these experiments, given the known requirement of HS PGs for successful exit from pluripotency and differentiation into neural precursor cells (40). Upon treatment of HS-deficient Ext1^−/−^ mESCs with cholPEGNTA, followed by a poly-His-tagged GFP (GFPHis_6_), we observed dose-dependent increases in cellular fluorescence in flow cytometry analysis (**Fig. S18**). Fluorescence was dependent upon the concentrations of cholPEGNTA and GFPHis_6_, and maximal fluorescence was achieved at higher doses, indicating that the presentation of molecules (~650,000 molecules of equivalent soluble fluorochrome; **Fig. S18**) using this strategy could be saturated.

**Figure 4.**
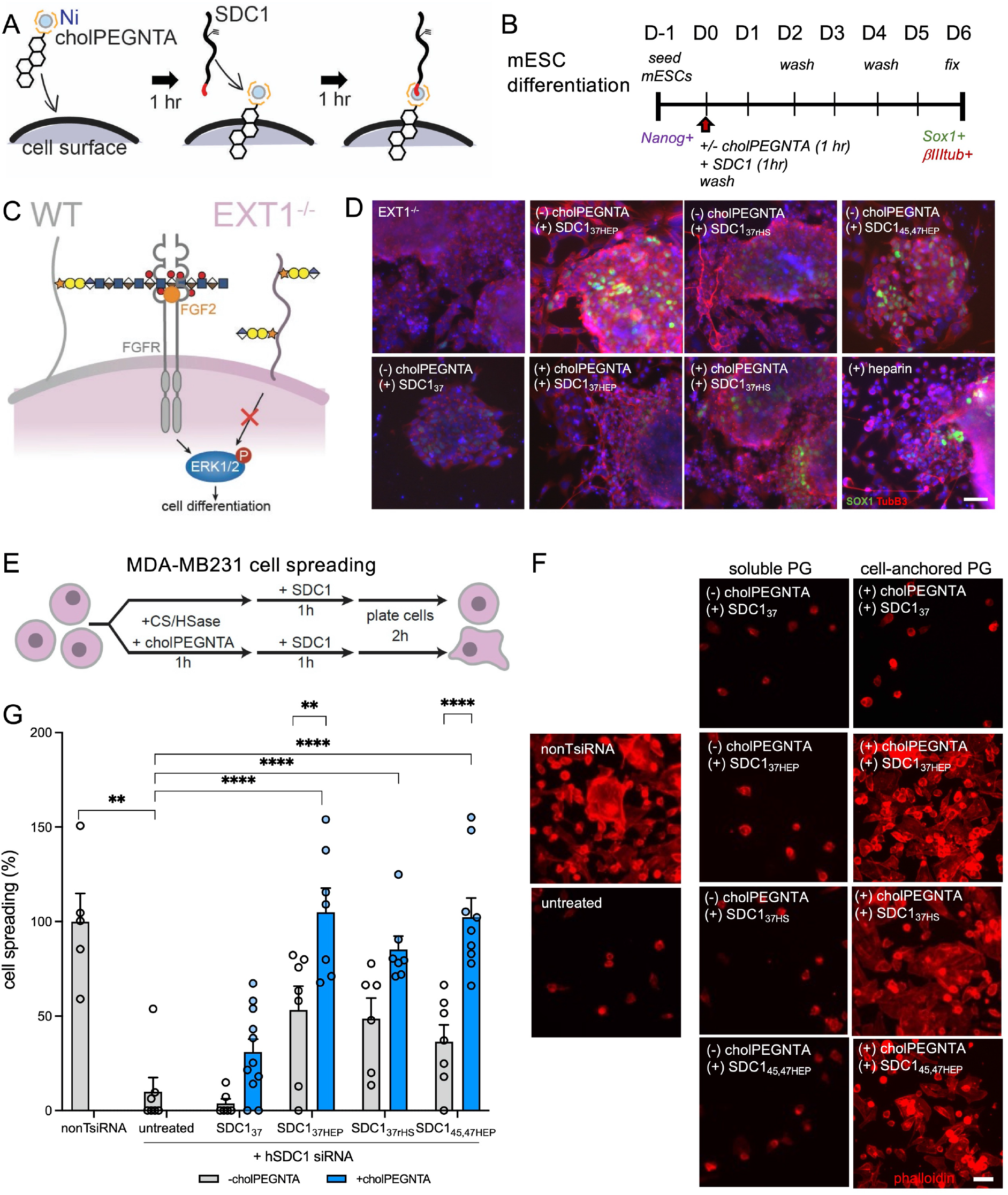
Cell surface engineering with edited SDC1 ectodomains and their effects on mESC differentiation and cancer cell spreading. (**A**) A two-step complexation is used to present engineered SDC1 ectodomain variants onto cell surfaces. Live cells are first incubated with cholPEGNTA (1hr, 37°C), followed by Ni(OAc)_2_ and the SDC1 ectodomain equipped with a poly-His tag (red tail; 1 hr, 37°C). Excess material is washed off at each step. We did not observe significant toxicity in mESCs with the introduction of any of these materials. (**B**) A six-day differentiation protocol for mouse embryonic stem cells, illustrating the loss of pluripotency marker Nanog and increased SOX1 expression, indicative of neuroectoderm differentiation. (**C**) In contrast to wild-type (WT) cells, mouse embryonic stem cells deficient in heparan sulfate (Ext1^−/−^) fail to differentiate due to their reduced ability to bind FGF2 (orange) effectively. (**D**) GAG-conjugated SDC1 ectodomains (2 μM) can rescue differentiation of Ext1^−/−^mESCs, as evidenced by expression of neural precursor markers Sox1 (green) and Tubb3 (red). Scale bar: 50 μm (**E**) Workflow for MDA-MB231 cancer cell remodeling spreading assay on vitronectin-coated surfaces. (**F**) Representative fluorescence microscopy images of SDC1-knockdown MDA-MB231 cells following incubation with or without cholPEGNTA (10 μM, 1 hr, 37°C) and with the engineered SDC1 ectodomains, and stained for phalloidin (red). Cells deficient in SDC1 fail to spread on vitronectin surfaces unless supplemented with membrane-anchored SDC1 conjugated to GAGs. Scale bar: 50 μm (**G**) Quantification of cell spreading on vitronectin surfaces. Bar graphs represent means and error bars represent standard deviations. For each condition, n >5 images. One-way ANOVA **** *p* <0.0001, ** p: 0.0017.

Using these optimized conditions, we subjected Ext1^−/−^ mESCs to a six-day adherent differentiation protocol (**Fig. 4B**) (41), where wild-type mESCs gradually lose expression of pluripotency markers and differentiate into SOX1- and βIII-tubulin (TubB3)-positive neural precursor cells. On the other hand, Ext1^−/−^ mESCs retain their pluripotent states unless they are supplemented with HS-presenting PGs or exogenous heparin (**Fig. 4C**). We incubated Ext1^−/−^ mESCs with various soluble (-cholPEGNTA) or membrane-anchored (+cholPEGNTA) SDC1 ectodomains for 1 hour only at the onset of the differentiation protocol (at D0) and analyzed the cells for expression of markers of ectodermal lineage and pluripotency after six days (D6, **Fig. 4D, Fig. S19**). As expected, we observe increased SOX1 expression in cells treated with our GAG-conjugated SDC1 constructs compared to untreated and SDC1_37_-treated Ext1^−/−^ mESCs. This is concurrent with a marked decrease in the expression of the pluripotency marker Nanog, indicating successful exit from pluripotency in cells treated with replete SDC1 constructs (**Fig**. **S19)**. Similar results were obtained from a three-dimensional embryoid body differentiation format (**Fig. S20**), which mimics hallmarks of early embryonic development. Remarkably, the extent of differentiation with our conjugates was similar to mESCs treated with soluble heparin despite its prolonged treatment, (heparin is present in culture from D0-D2). In contrast, mESCs were treated with our soluble engineered ectodomains for only 1 hour on D0, with excess, unbound material washed away. The dependence on GAG-conjugation for differentiation reinforces their requirement for binding FGF2 to trigger ERK1/2 phosphorylation (**Fig. S21**). FGF2 is known to initiate differentiation in mESCs by forming ternary complexes with FGFR and HSPGs, such as SDC1. Ext1^−/−^ mESCs stain positively for SDC1, which is presumably conjugated to CS (**Fig. S22**), yet these cells remain pluripotent when void of external triggers. These observations overall highlight the requirement for HS GAGs for mESC differentiation, with less of a requirement for membrane presentation.

### Cell surface engineering and cancer cell spreading

Syndecans are implicated the attachment and spreading of cells on the extracellular matrix (42). Thus, we also evaluated whether our SDC1 constructs could also rescue the spreading of human mammary cancer cells (MDA-MB231) in an established cell spreading assay (**Fig. 4E**) (42). Using SDC1-knockdown and GAG-digested cells (**Fig. S23**), we observed that all soluble forms of the SDC1_37_ ectodomain failed to rescue cell spreading on vitronectin matrices, and that only membrane-anchored GAG-conjugated variants were able to cause cell spreading (**Fig. 4F, Fig. 4G**). Similar results were obtained for SDC1-knockout cells obtained via CRISPR-Cas9 (**Fig. S24**). The most significant rescue was observed with the divalent heparin SDC1_45,47HEP_, followed by the monovalent SDC1_37HEP_. Additionally, the preference for heparin conjugates over HS is observed when comparing SDC1_37HEP_ vs SDC1_37HS_. The requirements of membrane-anchoring and GAGs are consistent with the significant affinity of α_v_β_3_ integrin for heparin (**Fig. 3F**), (34) and they are concurrent with previous observations by Beauvais et al., highlighting the dependence of membrane-bound SDC1 in vitronectin-mediated cell spreading (42). Because cell surface α_v_β_3_ binds both cell surface HS-conjugated SDC1 and the immobilized vitronectin (with an apparent K_D_ < 10 nM), it is likely that cell spreading is mediated by α_v_β_3_-SDC1-vitronectin ternary complexes and fine-tuned by *cis* and *trans*-occurring interactions, respectively. While deglycosylated SDC1 may also facilitate cell spreading via other protein-protein interactions or ligands, (42) due to the large enhancement in binding afforded by GAG-conjugated SDC1s, it is likely that the relevant biological mechanism is GAG-mediated.

### Proteoglycan editing platform is compatible with live cell proximity tagging

In addition to using our engineered SDC1 ectodomains in phenotypic assays, we also envisioned using the APEX2-tagged PGs to map the interactomes for PGs in live cells. PGs, such as SDC1, possess many binding partners as a consequence of their multi-component nature (43), which may vary as a result of differential glycosylation or cell surface localization. Understanding the structural requirements for PG interactors can aid in assigning mechanisms of action for their biological functions. As an initial proof of concept study, we used A-SDC1_45,47_ and its heparin conjugate A-SDC1_45,47HEP_ in a proximity tagging experiment to capture their interactors in live mESCs (**Fig. 5A**). Proximity tagging has gained recognition as a powerful approach to capture the interacting proteins for a given protein bait. In this approach, the APEX2 peroxidase enzyme catalyzes the formation of short-lived (< 1 ms) and proximal (<20 nm radius) biotinyl radicals that covalently react with proteins interactors for the bait protein. As such, the interacting proteins are biotinylated while the cells are alive, without disrupting their endogenous localization. The biotinylated proteins can then be probed using fluorophore-conjugated streptavidin and imaged using fluorescence microscopy or enriched for mass spectrometry-based identification. The APEX2 approach is especially apt for glycan-protein interactions that are heavily influenced by three-dimensional presentations of ligands, and which may be dynamic or weak (44).

**Figure 5.**
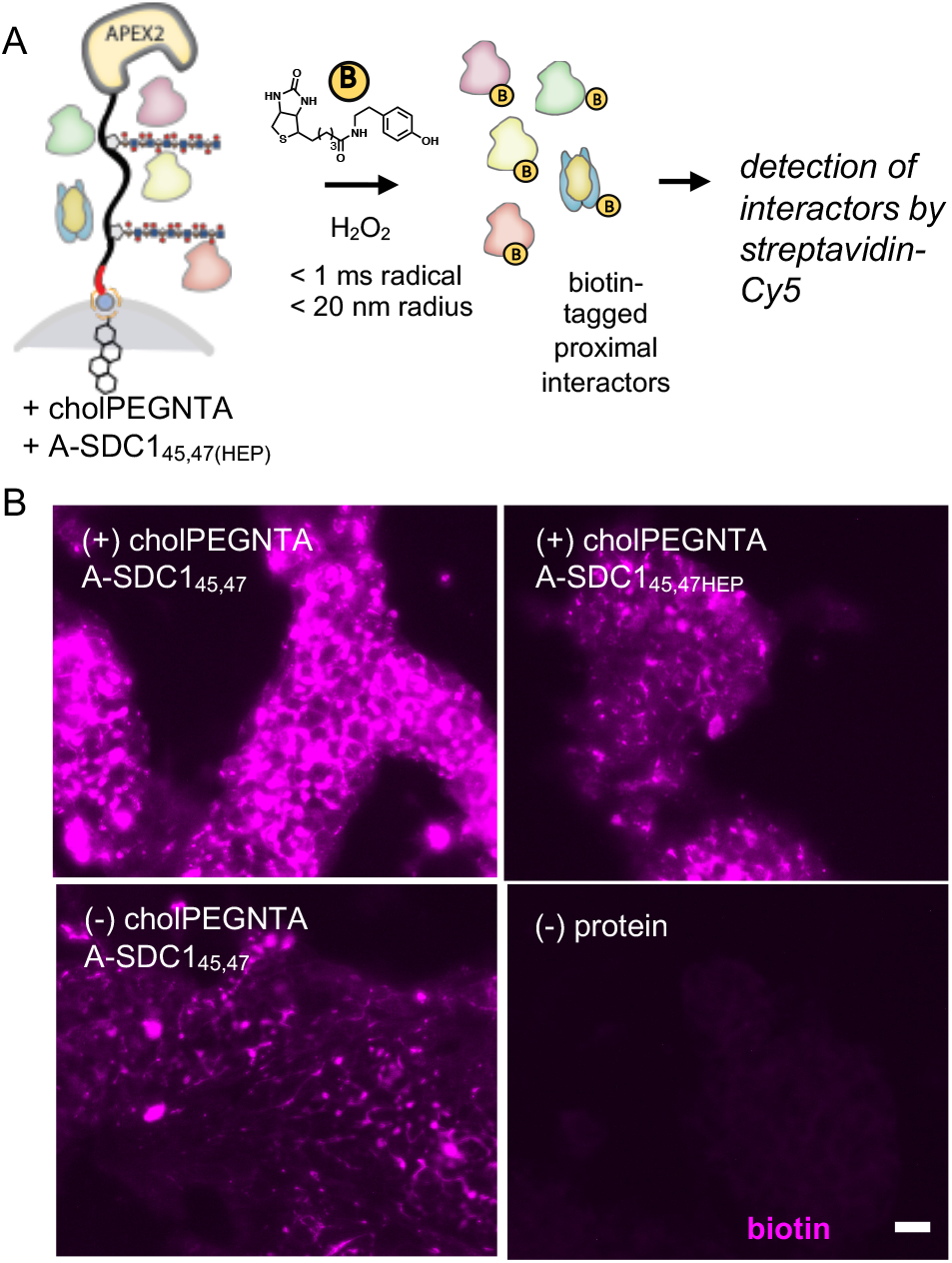
Proximity tagging with A-SDC1. (**A**) Live mESCs were incubated with A-SDC14_5,47_ or A-SDC1_45,47HEP_ (2 μM, 1 hr), with or without pre-treatment with cholPEGNTA (10 μM, 1 hr), to generate cell surface-anchored or soluble SDC1 ectodomains, respectively. Following treatment with biotin-phenol (yellow circle, 0.5 mM, 30 min) and a brief incubation with H_2_O_2_ (100 mM, 1 min), the APEX2 peroxidase catalyzes the formation of reactive biotin radicals that are short-lived (< 1 ms) and react with proteins located proximally (< 20 nm) from the complex. Biotinylated interactors are detected using Cy5-streptavidin. (**B**) Wild-type mESCs subjected to the proximity tagging protocol demonstrate robust fluorescence signals from biotinylated interactors. The signals are mostly located at the cell surface, compared to the non-treated control. Membrane anchoring and glycosylation appear to regulate the extent of fluorescence, and thus the interactomes, of SDC1 fusion proteins.

We qualitatively compared the differential interactomes of A-SDC1_45,47_ when it is presented as a soluble or a cell-surface anchored molecule. Following incubation with cholPEGNTA, mESCs were similarly incubated with Ni(OAc)_2_ and the A-SDC1_45,47_ ectodomains. We observed significant dose-dependent (**Fig. S25**) biotinylation of mESCs following proximity tagging (**Fig. 5B**). The majority of the signals arose from the cell surface, in both WT and Ext1^−/−^ mESCs (**Fig. S26**) consistent with the known localization of SDC1. Interestingly, we also observed varying intensities of biotinylation depending on the presence of the GAG chain and the presentation of the fusion protein, with highest signals were observed from membrane-anchored, deglycosylated A-SDC1_45,47_. The detection of fluorescence in mESCs treated with soluble versions of A-SDC1_45,47_ was expected, given that SDC1 is known to interact with cell surface ligands, such as integrins (**Fig. 3D**). While we have yet to assign the differential interactomes resulting from changes in PG architecture, we demonstrate that fusion proteins of our engineered PGs have tremendous utility and promise to fine-tune our understanding of native PG interactions in various contexts with minimal perturbation.

## Discussion

PGs are important information-dense molecules that reside at the cell surface. Because glycosylation is not directly controlled by the genetic code, PGs naturally exist as molecular polymorphs of various GAG motifs attached to a single core protein. It is widely speculated that the molecular heterogeneity found within PGs is important as it largely contributes to their functional diversity, although this remains poorly understood due to the challenges of studying PG architecture. To begin systematically dissecting the effects of PG molecular heterogeneity toward function, we have assembled a holistic platform to chemically edit PGs using an integrated approach that merges unnatural amino acid incorporation, metabolic oligosaccharide engineering, and cell surface engineering. Starting with the core proteins of SDCs 1-4, we incorporated unnatural amino acids at sites of native GAG glycosylation for bioorthogonal conjugation (**Fig. 1**). Notably, we recapitulated the endogenous multi-glycosylated nature of SDC1, which is known to be modified by HS at two sites.

GAGs are among the most complex and ubiquitous forms of glycan chains. While much heroic work has focused on the total synthesis of short GAG oligomers and proteoglycan-mimetic glycoconjugates, thus far, these products remain limited in their ability to capture the sophistication of PG architectures. We demonstrate the use of a metabolic oligosaccharide engineering approach to append bioorthogonal labels in GAGs, which are traceable using CuAAC (**Fig. 2**). Owing to the common tetrasaccharide linker for both HS and CS GAGs, the azidoxyloside **1** generates azide-modified versions of both. Using selective enzymatic digestion, we are also able to generate homogenous GAGs (i.e. HS vs CS) despite the fact they are produced from cells as a mixture. These homogeneous GAG samples will be especially valuable toward understanding the functional relevance of PGs that are present as various hybrids of HS or CS. While we have yet to fully map the intermediate metabolic conversions the compound undergoes once inside the cell, it likely undergoes similar mechanisms as other xylosides (28, 31, 45).

We demonstrate here that CuAAC, a widely used bioorthogonal reaction, can be used to conjugate a highly polyanionic GAG molecule to one or two sites of an equally large PG core protein (**Fig. 3**). Owing to its bioorthogonal nature, this conjugation method is relatively inert, and it is compatible with heme-containing proteins, such as the APEX2 peroxidase. Although the GAGs produced by compound **1** remain heterogeneous in sizes, they retain the macromolecular feature and domain-like arrangement of sulfation motifs of natural GAGs. In studying the binding interactions between the engineered SDC1 ectodomains, we uncovered that the divalent SDC1_45,47HEP_ binds more tightly toward α_v_β_3_ compared to the monovalent SDC1_37HEP_, which was functionally recapitulated in cell spreading. These observations could be attributed to potential synergistic or cooperative effects, although we cannot rule out the potential for positional importance, where specific positions among the three GAG sites of SDC1 can contribute more towards binding than others (25). While we have made significant strides to achieve defined yet native architectures for PGs, there remain opportunities to expand the repertoire of edited PGs using multiple UAAs (46) and other synthetic xylosides primers.

In contrast to standard genetic approaches, our platform allows the facile interrogation of the roles of specific GAG structures within the context of the native PG core protein, and it permits us to tune the presentation of PG ectodomains. Altering the presence and composition of GAG chains via genetic means can disproportionately cause higher expression levels of certain SDC1 mutants or affect the glycosylation of other GAG sites (47), complicating the analysis of the effects of GAG deletions (25). Our platform not only provides a means to generate defined PG ectodomains, but coupled to cell surface engineering strategies, it greatly facilitates the study of native PG glycoconjugate interactions functions at surfaces of live cells, especially in those that are not readily amenable to genetic manipulation (e.g. primary cells, stem cells). The ability to perform these structure-activity studies is crucial, as many organisms and proteins that interact with HS GAGs *in vitro* have been observed to not interact to native HSPGs (48). These differences could be due to additional binding contributions of the core protein itself, the differing fine structure of HS among tissues, or the absence of the multivalent architectural arrangement naturally present in PGs. To our knowledge, this work represents the most advanced effort toward the generation of full PG glycoconjugates with defined GAG chains.

We have demonstrated the utility of our platform in two functional phenotypic assays, mESC differentiation (**Fig. 4D**) and cancer cell spreading (**Fig. 4G**). Both systems illustrate the requirement for the correct GAG compositions in order to exhibit positive effects, yet it is only with vitronectin-mediated cancer cell spreading that we observe an obligate requirement for cell surface anchoring. These observations could be due to the fact that the FGF2 is also present as a soluble molecule in mESC differentiation, whereas in cancer cell spreading, both vitronectin and the α_v_β_3_ integrin are anchored either as a substrate or on the cell surface, such that ternary complex formation is much more difficult to achieve with soluble SDC1. However, we cannot rule out the possibility that there may be different extents of ternary complex formation in these two systems, and an auto-inhibitory effect may be achieved with soluble SDC1 at higher concentrations (49).

The ability to tag interactors in live mESCs by combining cell surface engineering with proximity tagging open up new opportunities to study PG interactions. Notably, SDC1 alone exhibits over 100 interactors in different contexts. (43) Our cell surface engineering approach is applicable to any cell with a phospholipid membrane, and it is especially useful for mESCs, which are generally recalcitrant towards transient genetic manipulations. As such, stem cells are not amenable to structure-activity relationship studies of PGs that require knock-in/knockdown of individual components. Studying PG biology at the cell surface, where they are free to interact with their ligands and move along the plane of the membrane, may also allow focal concentration of the PGs to enact their corresponding functions. SDC1 is an important PG that mediates myriad complex biological functions. When anchored at the cell surface, it can serve as a functional receptor for microbes (50) and cytokines, and promote the assembly of extracellular matrix components (51). Interestingly, when it is processed by matrix metalloproteinases or sheddases, the soluble ectodomains can carry out competing functions to membrane-anchored SDC1 (52). Thus, we expect that further inquiry into these biological contexts may benefit from our platform. While there are 16 classically known PGs, of which we have made progress on four, recent work has identified that other proteins not conventionally designated as PGs, can bear GAG chains and carry out important biological functions (53, 54). These discoveries therefore create even further opportunities for the use of glyco-engineering platforms, like ours, to aid in our understanding of the biology of protein glycoconjugates.

## Conclusion

In summary, we have generated a modular platform that permits the tailored assembly of proteoglycan ectodomains, replete with sequence-specific core proteins, defined GAGs, and the ability to present them onto cell surfaces. The capacity to site-specifically incorporate a growing number of functionalized unnatural amino acids into PG backbones provides new opportunities to study their structural conformations, as well as chemoselectively conjugate additional moieties (55, 56). The efficient metabolic oligosaccharide engineering approach we used to generate working amounts of azide-modified GAGs is compatible with multiple cell types, and sulfation-mutant cell lines will greatly facilitate studies on native GAGs beyond heparin (57–59). The GAG-conjugated SDC1 proteoglycans used here represent the most defined semi-synthetic materials (60) to date, and they capture many functional facets of PG ectodomains. We have demonstrated that the systematic variation of GAG composition and multivalency on SDC1 core proteins can yield refined insights into structure-function relationships, as it relates to mESC differentiation and cancer cell spreading. These materials may also serve beneficial for controlled localized delivery of GAG-binding proteins (60). The process for producing these conjugates is relatively simple - over the course of these studies, we have produced sub-milligram quantities of the most complex material in the series, SDC1_rGAG_. Lipid-based cell surface engineering and radical-based proximity tagging approaches are readily assimilated into the engineered proteoglycans, and we anticipate that these materials and this platform will be further utilized to dismantle the complexity of proteoglycans in many other biological contexts.

## Methods

### Materials and reagents

Unless otherwise noted, chemical reagents were used as purchased and obtained from commercial vendors. Recombinant human α_v_β_3_ (R&D Biosystems # 3050-AV-050) is composed of the α_v_ (Phe31-Val992, Accession # NP_002201) and the β_3_ (Gly27-Asp718, Accession # AAA52589) subunits. Recombinant human FGF2 (NovusBio # NBP2-76301) is composed of 154 amino acids (Accession Number: P09038). Wild-type mESCs and MDA-MB231 cells were purchased from ATCC, and Ext1^−/−^ mESCs were a gift from Kamil Godula, Catherine Merry, and Jeffrey Esko.

### Expression of pPY-modified PG ectodomains

The syndecan ectodomain cDNA sequences were derived from EMbL, mutated as desired, codon-optimized for bacterial expression, synthesized (Genscript, NJ, USA), and cloned into the NheI/NotI sites of the pET-28a(+) vector with kanamycin resistance. Rosetta 2 (DE3) competent (Novagen) bacterial cells were co-transformed with the SDC-containing plasmid and pULTRA-CNF (Addgene # 48215, a gift from Peter Schultz) [14]. Following spectinomycin and kanamycin-mediated selection, single colonies were picked and expanded in LB media containing the same antibiotics and cultured until OD_600_ ~ 0.6-0.8. IPTG and the pPY unnatural amino acid (400 mg/L) were then added, and the culture was further incubated overnight. The cells were then harvested, lysed, and purified using FPLC systems equipped with nickel-NTA columns to harvest the SDC1 ectodomains. Following elution with imidazole and subsequent purification by anion exchange or size exclusion chromatography, proteins were stored in PBS with 25% glycerol, flash frozen in methanol/dry ice and stored at −80 °C.

### Chemical synthesis of azidoxyloside 1 (Fig. 2B)

See supporting information for full chemical synthesis and purification.

### Molecular docking of azidoxyloside 1 with β4GalT7

Compound **1** was docked into the active site of β4GalT7 in complex with UDP-galactose (PDB ID 4M4K) (61) using Autodock Vina (62).

### Production of recombinant GAGs by azidoxyloside priming

CHO-K1 cells were seeded on 15 cm^2^ dishes overnight and incubated with **1** (400 μM, 48-72 hr) in basal DMEM media. After the desired incubation period, the conditioned media (CM) was collected, lyophilized, and purified using manual DEAE columns. Dialysis with water was performed to remove excess salt.

### Disaccharide analysis and quantification of azide-primed GAGs

Harvested GAGs were digested overnight with 2.5 U/mL heparinases I, II, III (Sigma) or 0.5 U/mL chondroitinase ABC (Sigma). Disaccharides were isolated with a 3000 MWCO filter (Thermo) and labeled with anthranilamide (63). Labeled disaccharides were analyzed with a Propac PA1 column (ThermoFisher Scientific) on an Ultimate 3000 UHPLC with fluorescence detection (57). To gauge the extent of azide incorporation, GAGs were reacted with sulfo-DBCO-biotin (Sigma) (6.7 mM, 20 h, 37 C, PBS). Excess DBCO-biotin was removed with size-filtration, and GAGs were incubated with streptavidin agarose (Sigma). Unbound GAGs were separated from GAGs captured on the streptavidin beads, and both fractions were treated with either heparinase or chondroitinase. Disaccharides were labeled and analyzed and proportions of azide-labeled GAGs were measured by comparing fluorescence signals for bead-captured vs. unbound fractions.

### CuAAC conjugations

CuAAC conjugations (60) were performed in PBS (pH 6-8) with the proteoglycan (25 μM), azide-GAG (900 μM), aminoguanidine hydrochloride (5.0 mM), CuSO_4_ (320 μM), Tris(3-hydroxypropyltriazolylmethyl)amine (THPTA; 1.6 mM), and sodium ascorbate (21 mM), 37 °C for 2 hr. Conjugates were purified using weak anion exchange DEAE chromatography (WAX-10 4×250 mm column, 0 to 400 mM NaCl, 15 min, then 1.75 M NaCl to elute). DEAE eluate was loaded onto a Ni-NTA column, and washed to remove unconjugated azide-GAGs (1 M NaCl tris buffer). The reaction product was eluted with imidazole, desalted, and concentrated prior to analysis or subsequent use.

### ELISA with immobilized syndecans

Each SDC1 construct was immobilized in duplicate using pH 9.6 carbonate buffer onto 96-well high-binding plates (4°C, overnight, rocking, 50 mL, 10 mg/mL) (Thermo Maxisorp plates, 439454). Following blocking (2% BSA/PBST, 100 mL) and washing 3X with PBST (100 mL), wells were incubated with titrations of biotinylated FGF2 or α_v_β_3_ (PBST, 4°C, overnight, rocking, 50 mL). Following 3X washes with PBST, wells were probed with HRP-streptavidin (Biolegend 405210, 1000:1) or biotinylated anti-a α_v_β_3_ antibody (Biolegend 304412, 1:1000) (RT, 1 hr, rocking, 100 mL) followed by washing and HRP-streptavidin (RT, 1 hr, rocking). Wells were then washed 3X with PBST and incubated with a TMB substrate (BioFX, 100 mL) until a solid blue color develops, and the reaction was quenched with sulfuric acid (2N, 100 mL). Absorbance values at 450 nm was read and subtracted from blank (no FGF2 or α_v_β_3_) wells, normalized to 100%, and fitted into a non-linear logarithmic curve (Prism).

### Cell surface engineering

Cells were washed with DPBS and incubated with cholPEGNTA (Nanocs # PG2-CSNT-2k; 10 μM, 1 hr, 37°C) After washing twice with DPBS, cells were incubated with SDC1 constructs (pre-centrifuged at 14,000 xg, 10 min. to exclude potential aggregates in KO-DMEM media (1 hr, 37°C). Cells were then washed 2X with DPBS and re-suspended in N2B27 media for 6 days to differentiate. For neuronal differentiation, cells were cultured in differentiation (N2B27) media for 6 days (**Fig. 4B**). For the cell spreading assay, cells were plated onto vitronectin-coated plates (**Fig. 4E**).

### Neuronal differentiation of mouse embryonic stem cells (mESCs)

For adherent mESC differentiation (64), cells were seeded overnight (37°C, 5% CO_2_, ~18 hr) in maintenance media (MM) at a density of 1×10^4^ cells/cm^2^ in 24-well gelatin-coated 24-well plates. The following day (D0), mESCs were washed 1X with PBS and the media was replaced with N2B27 neural differentiation media. Subsequent washes and media replacements were performed at D2, D4, and the cells were fixed at D6, stained for various markers, and imaged using fluorescence microscopy.

### Cell Spreading Assay

SDC1-knockdown GAG-digested MDA-MB-231 cells harvested using a non-enzymatic cell dissociation buffer were seeded (5×10^4^ cells per well) in 24-well plates pre-coated with poly-D-lysine and vitronectin (2 hr at 37°C, 5% CO_2_). After washing 1X with PBS and fixation, cell spreading was visualized using rhodamine-conjugated phalloidin and imaged using fluorescence microscopy.

### Live cell proximity labeling

Proximity tagging was adapted from previously published procedures (44, 65). mESCs seeded on gelatin were treated with cholPEGNTA (10 μM, 1 hr, 37°C) and further incubated with APEX2 fused proteoglycans (10 μM, 1 hr, 37°C) after washing 2X DPBS. After further washes, cells were incubated with biotin phenol (500μM in KO-DMEM, 30 mins at 37°C), followed by H_2_O_2_ (1 mM, 1 min, RT). The reaction was stopped with 3X washes of quenching solution (5 mM Trolox, 10 mM sodium ascorbate, and 10 mM sodium azide in DPBS). mESCs were then fixed, blocked, probed for biotinylated interactors by Cy5-streptavidin, and imaged by fluorescence microscopy.

## Author contributions

T.O., M.C., T.N.S., and M.L.H. conceived and designed the research. T.O. performed molecular docking simulations. T.O. and X. Y. expressed engineered proteoglycans and performed bioconjugation reactions. T.N.S., N.B., and R.H. performed xyloside priming experiments and T.O., T.N.S., and N.B. performed GAG analysis. M.C. performed mESC differentiation, cell spreading assays, and proximity tagging experiments. X. Y. contributed to biological assays. T.O., M.C., T.N.S., and M.L.H contributed to manuscript preparation.

## Acknowledgements

We thank Drs. Sourav Chatterjee and Abdullah Hassan for their assistance in preparing the azidoxyloside compound. This work, T.N.S. (HD090292-S1), M.C., and M.L.H are supported by the NIH K99/R00 Pathway to Independence Award (R00-HD090292). M.L.H is grateful for the support and scientific counsel of Profs. Kamil Godula and Jeffrey Esko for this work.

