## Supporting Information for "Chemical Editing of Proteoglycan Architecture"

CONTENTS:

**Table S1.** List of reagents and sources 3-4
Syndecan ectodomain amino acid sequences 5

**BIOCHEMICAL AND SYNTHETIC METHODS** 6-11
Preparation of pPY unnatural amino acid
Expression of syndecan ectodomains with pPY amino acids
Mass spectrometry of intact $SDC1_{37}$.
MMP9/Sheddase treatment of $SDC1_{37}$.
Preparation of azidoxyloside **1.**
Preparation of azido-heparin or azido-CS.
Production, purification, and analysis of recombinant GAGs from CHO-K1 cells (**Scheme S1**).
Preparation of chondroitinase ABC and heparinase cocktail.
Enzymatic digestion of GAGs.
GAG disaccharide labeling.
GAG disachharide analysis.
CuAAC reactions.
HPLC monitoring of click reactions.
Biotinylation of FGF2 for ELISA

**BIOLOGICAL METHODS** 11-12
Flow cytometry following cell surface engineering
Neuronal differentiation of mouse embryonic stem cells
Immunofluorescence staining
FGF2 stimulation assay
Knockdown of SDC1 by siRNA or CRISPR/Cas9 in MDA-MB231 cells
Quantification of cell spreading

**ADDITIONAL DATA** 13-29
**Figure S1A-C.** Characterization of $SDC1_{37}$. 13
**Figure S2A-C.** Characterization of A-$SDC1_{45,47}$. 14
**Figure S3A-C**. SDS-PAGE analysis of $SDC2_{41,55,57}$, $SDC3_{80,82,89}$, and $SDC4_{44,62,64}$ 15
**Figure S4A-C.** Sheddase activities of engineered $SDC1_{37TAMRA}$. 16
**Figure S5**. Expression of $GPC1_{485,487,489}$.
**Figure S6**. $^1$H NMR analysis of azidoxyloside **1**. 17

**Figure S7.** Molecular docking of **1** or its non-azide derivative with β4GalT7.
**Figure S8A-B**. Characterization of recombinant GAGs isolated from CHO-K1 cells. 18
**Figure S9A-B.** Analysis of HS disaccharide compositions.
**Figure S10A-B.** Figure S10. Recombinant azide-primed GAGs recapitulate sulfation patterns of host GAGs. 19
**Figure S11.** Recombinant azide-primed CS are similar despite differing incubation times with **1.**
**Figure S12.** Azido-heparin mimics the composition of native heparin**.**
**Figure S13A-B.** $^{1}$H NMR (400 MHz, $D_2O$) analysis of azido-GAGs 20
**Figure S14A-E.** Additional data for $SDC1_{37}$ conjugated to azido-GAGs 21
**Figure S15.** A-$SDC1_{45,47}$ conjugation to azido-GAGs. 22
**Figure S16.** $SDC4_{44,462,64}$ conjugation with azido-heparin.
**Figure S17A-D.** Conjugation of pPY-containing GFP-Y to azidoGAGs**.** 23
**Table S1 and S2.** ELISA data for $\alpha_v\beta_3$ integrin or FGF2 binding to SDC1 ectodomains. 24
**Figure S18.** Cell surface engineering with GFP-$His_6$ as a model. 25
**Figure S19.** Differentiation of adherent mESCs at D6 of differentiation.
**Figure S20.** Embryoid body (EB) differentiation at D6. 26
**Figure S21.** FGF2 triggers the activation of the ERK pathway. 27
**Figure S22.** mESCs express SDC1.
**Figure S23.** MDA-MB231 cells treated with 100 or 200nM pooled dsRNAi exhibit reduced SDC1 expression
**Figure S24.** Rescue of cell spreading in $SDC1^{KO}$ cells..
**Figure S25.** Proximity tagging across different concentrations. 28
**Figure S26.** Proximity tagging in WT and $Ext1^{-/-}$ mESCs. 29
**References** 29

**Table S1. List of reagents and sources**

| Product | Manufacturer | Product # |
|---|---|---|
| Type A, Porcine Skin Gelatin Powder | Sigma | G-1890 |
| KO-DMEM | Gibco | 10829-018 |
| Gemcell FBS | Gemini Bioproducts | 100-500 |
| Leukemia inhibitory factor (LIF, ESGRO) | Millipore | ESG1107 |
| MEM NEAA (100X) | Gibco | 11140-035 |
| 2-mercaptoethanol (2-ME, 50mM) | Gibco | 31350-010 |
| Neurobasal | Gibco | A35829-01 |
| DMEM/F-12 | Gibco | 11320-033 |
| L-glutamine (200mM) | Gibco | 25030-024 |
| N-2 supplement (100X) | Gibco | 17502-048 |
| B-27 Plus supplement (50X) | Gibco | A35727-01 |
| Heparin | Iduron | Hep001 |
| Lab-Tek II chamber slide system (8-well) | Thermofisher | 154534 |
| Cholesterol-PEG-NTA | Nanocs | PG2-CSNT-3k |
| EDTA-free trypsin | Quality Biological | 118-086-721 |
| Non-enzymatic cell dissociation buffer | PeproTech | CPD-125 |
| Lipofectamine RNAiMAX | Invitrogen | 13778-150 |
| Vitronectin | PeproTech | 140-09-1mg |
| Recombinant human fibronectin fragment 2 | Sino Biological | 10314-H08H |
| GFPHis6 | Addgene | plasmid # |
| FGF2 | Novus | NBP2-76301 |
| AVB3 | ACRO | IT3-H52E3 |
| Trypsin-EDTA (0.05%) | Gibco | 25300-062 |
| Trypsin-EDTA (0.25%) | Gibco | 25200-056 |
| IPTG | Bioworld | 21530057 |
| Kanamycin | Fisher | BP906-5 |
| Spectinomycin | GoldBio | S-140-25 |
| Imidazole | Sigma-Aldrich | 12399 |
| Aminoguanidine hydrochloride | Acros | 368910250 |
| Copper sulfate | Acros | 42287-1000 |
| THPTA | TCI | T3171 |
| Sodium ascorbate | ChemImpex | 1436 |
| 5-TAMRA-azide | Lumiprobe | 37130 |
| Anthranilamide (2-AB) | Acros | 104900050 |
| 4-(bromomethyl)benzoate | Combiblocks | OR-0319 |
| Sodium azide | Fisher | BP9221 |
| DMF | Fisher | D119-1 |
| Deuterium Oxide | Aldrich | 151882 |
| DMSO-D6 | Aldrich | 256147 |
| Heparin Agarose beads | Sigma | H0404 |
| Sodium Acetate | Sigma | 791741 |
| Aniline | Sigma | 24284 |

| | | |
|---|---|---|
| DMSO | Fisher | D128-1 |
| Fluor-488-alkyne | Sigma | 761621 |
| Sulfo-DBCO-biotin | Sigma | 760706 |
| Heparinase I and IIII | Sigma | H3917 |
| Heparinase II | Sigma | H6512 |
| TMB HRP substrate | BioFX | TMBW-1000- |
| Streptavidin agarose resin | Thermo | 20353 |
| Chondroitinase ABC | Sigma-Aldrich | C2905 |
| Sodium cyanoborohydride | Sigma-Aldrich | 156159 |
| MMP-9 | R&D Biosystems | 909-MM |
| 4-Aminophenylmercuric acetate | Sigma-Aldrich | A9563 |
| ADAM-TS1 | R&D Biosystems | 2197-AD |
| Heparin disaccharide standard mix | Galen | HD Mix |

| **Primary antibody** | **Manufacturer** | **Product #** |
|---|---|---|
| SOX1 | Cell Signalling | 4194S |
| Nestin (clone 4D4) | Invitrogen | 14-5843-82 |
| Nanog (clone D2A3) | Cell Signalling | 8822T |
| B-tubulin III (TubB3) | Proteintech | 66240-1-1g |
| SDC1 (clone 281-2) | BioLegend | 142502 |
| SDC1 (clone MI-15) | BioLegend | 356502 |
| Rhodamine-phalloidin | Biotium | 50-196-4057 |
| ERK1/2 | Cell Signalling | 4695S |
| Phospho-ERK1/2 | Cell Signalling | 4370S |
| Alpha-tubulin | Abcam | ab7750 |
| human CD51/61 | Biolegend | 304412 |

| **Secondary antibody** | **Manufacturer** | **Product #** |
|---|---|---|
| Donkey anti-mouse IgG AF647 | Invitrogen | 1984047 |
| Donkey anti-mouse AF555 | Invitrogen | A31570 |
| Goat anti-rabbit IgG AF647 | Invitrogen | A32733 |
| Streptavidin-Cy5 | Southern Biotech | 7100-15 |
| Streptavidin-HRP | Biolegend | 405210 |
| Streptavidin-HRP | Biolegend | 405210 |
| Goat anti-rabbit IgG HRP | Abcam | Ab6721 |
| Goat anti-mouse IgG HRP | Invitrogen | G21040 |

**Proteoglycan ectodomain sequences**

**Syndecan-1 (18-252, wild-type)**

```
MQPALPQIVA VNVPPEDQDG SGDDSDNFSG SGTGALPDTL SRQTPSTWKD VWLLTATPTA PEPTSSNTET
AFTSVLPAGE KPEEGEPVLH VEAEPGFTAR DKEKEVTTRP RETVQLPITQ RASTVRVTTA QAAVTSHPHG
GMQPGLHETS APTAPGQPDH QPPRVEGGGT SVIKEVVEDG TANQLPAGEG SGEQDFTFET SGENTAVAAV
EPGLRNQPPV DEGATGASQS LLDRKEHHHH HH
```

SG = CS site
N = N-glycosylation site
SG = HS site
SS = MMP-9 cleavage site
L…E = interaction site b/w AVB3
pI = 4.71
Calculated Avg MW = 25422.56 Da

**Syndecan-2 (19-145, wild-type)**

```
METRTELTSD KDMYLDNSSI EEASGVYPID DDDYSSASGS GADEDIESPV LTTSQLIPRI PLTSAASPKV
ETMTLKTQSI TPAQTESPEE TDKEEVDISE AEEKLGPAIK STDVYTEKHS DNLFKRTEHH HHHHHHH
```

SG = HS site
pI = 4.70
Calculated Avg MW = 15394.59 Da

**Syndecan-3 (45-384, wild-type)**

```
MAQRWRNENF ERPVDLEGSG DDDSFPDDEL DDLYSGSGSG YFEQESGLET AMRFIPDMAL AAPTAPAMLP
TTVIQPVDTP FEELLSEHPS PEPVTSPPLV TEVTEVVEES SQKATTISTT TSTTAATTTG APTMATAPAT
AATTAPSTPE APPATATVAD VRTTGIQGML PLPLTTAATA KITTPAAPSP PTTVATLDTE APTPRLVNTA
TSRPRALPRP VTTQEPDVAE RSTLPLGTTA PGPTEMAQTP TPESLLTTIQ DEPEVPVSGG PSGDFELQEE
TTQPDTANEV VAVEGAAAKP SPPLGTLPKG ARPGPGLHDN AIDSGSSAAQ LPQKSILERK EVHHHHHHHH
HH
```

SG = CS site
SG = HS site
pI = 4.52
Calculated Avg MW = 36718.50 Da

**Syndecan-4 (24-145, wild-type)**

```
MESIRETEVI DPQDLLEGRY FSGALPDDED AGGSDDFELS GSGDLDDTEE PRPFPEVIEP LVPLDNHIPE
NAQPGIRVPS EPKELEENEV IPKRAPSDVG DDMSNKVSMS STAQGSNIFE RTEHHHHHHH HHH
```

SG = HS site
pI = 4.49
Calculated Avg MW = 14855.96 Da

**A-SDC1 (wt, residues 45 and 47 are serine)**

```
MGKSYPTVSA DYQDAVEKAK KKLRGFIAEK RCAPLMLRLA FHSAGTFDKG TKTGGPFGTI KHPAELAHSA
NNGLDIAVRL LEPLKAEFPI LSYADFYQLA GVVAVEVTGG PKVPFHPGRE DKPEPPPEGR LPDPTKGSDH
LRDVFGKAMG LTDQDIVALS GGHTIGAAHK ERSGFEGPWT SNPLIFDNSY FTELLSGEKE GLLQLPSDKA
LLSDPVFRPL VDKYAADEDA FFADYAEAHQ KLSELGFADA GSGGGGSENL YFQGMQPALP QIVAVNVPPE
DQDGSGDDSD NFSGSGTGAL PDTLSRQTPS TWKDVWLLTA TPTAPEPTSS NTETAFTSVL PAGEKPEEGE
PVLHVEAEPG FTARDKEKEV TTRPRETVQL PITQRASTVR VTTAQAAVTS HPHGGMQPGL HETSAPTAPG
QPDHQPPRVE GGGTSVIKEV VEDGTANQLP AGEGSGEQDF TFETSGENTA VAAVEPGLRN QPPVDEGATG
ASQSLLDRKE HHHHHHHHHH
```

Theoretical pI: 5.12
MW (Da): 54288.99 (average mass)

### BIOCHEMICAL AND SYNTHETIC METHODS

**Preparation of pPY unnatural amino acid.** The synthesis of pPY was completed in three steps starting from commercially available *N*-Boc tyrosine, following the procedures outlined by Deiters et al. [1] Briefly, *N-tert*-butoxycarbonyltyrosine (2 g, 7 mmol, 1 eq.) in (15 mL) DMF was slowly reacted with propargyl bromide (2.1 mL, 21 mmol, 3 eq., 80% toluene) overnight at RT. Following extraction with water and $Et_2O$,the organic layers were concentrated to yield the di-esterified product, a yellow oil, which was directly used in the next step. To a cooled (0 °C) solution of the crude material in methanol was slowly added a solution of 5 M HCl/MeOH until complete product formation. The solution was allowed to warm to room temperature and volatiles were removed to yield the propargyl ester as a yellow solid. The resulting product was dissolved in a mixture of 2 M NaOH and MeOH for 2 hr, RT. The pH was adjusted to 7 and the mixture was kept at 4 °C overnight. The resulting precipitate was washed with ice-cold water and dried *in vacuo*.

**Expression of syndecan ectodomains with unnatural amino acids.** Bacterial expression of syndecan ectodomains incorporating *p*-propargyl tyrosine began with co-transformation of Rosetta 2 (DE3) competent cells (Novagen). The syndecan ectodomain was encoded in a pET28a vector, was codon-optimized (Genscript) for expression in *E. Coli* and contained the TAG codon at the site of unnatural amino acid incorporation. The tRNA and engineered tRNA synthetase were in the pULTRA-CNF plasmid, a gift from Peter Schultz (Addgene plasmid # 48215 ; http://n2t.net/addgene:48215 ; RRID:Addgene_48215) [2]. Tubes containing 25 µL of competent cells were incubated with 200 ng of each plasmid on ice for 10 minutes, submerged in a water bath at 42 °C for 30 seconds, followed by 2 minutes on ice. At this point 100 µL of SOC media was added, and the cells were incubated at 37 °C for one hour, then selected by growing overnight on agar with 50 µg/mL each of spectinomycin and kanamycin. Single colonies were picked, grown for 18 hr at 37 °C in 4 mL of Luria-Bertani media with 50 µg/mL of spectinomycin and 50 µg/mL kanamycin, which was diluted with an equal volume of 50% glycerol and stored at -80 °C. This glycerol stock was used to seed overnight 4 mL cultures supplemented with antibiotics, and 2 mL of culture was added into 1 L of Luria-Bertami media with 50 µg/mL each of spectinomycin and kanamycin, in a 4 L flask. The media was shaken at 225 rpm at 37 °C until the $OD_{600}$ ~ 0.6 to 0.8. IPTG (1 mM) was then added, 13.3 mL of pPY (30 mg/mL in 250 mM NaOH with 15% DMSO) was added to a concentration of 400 mg/L, and an equal volume (13.3 mL) of 10X PBS (Fisher BP39920) solution was added as a buffer. The culture was shaken at 225 rpm at 30 °C for 18 hr. Cells were harvested by centrifuging for 20 minutes at 3600 x *g* at 4 °C, and subsequently washed with ice-cold PBS. Cells were resuspended in 20 mL of 3X concentrated PBS buffer (Fisher BP39920) with 25 mM of imidazole at pH 7.5 containing 1 mM phenylmethylsulfonyl fluoride (PMSF) and 5 µg/mL each of leupeptin (EMD, 108975), aprotinin (GoldBio A-655-25), and pepstatin A (Thermo, J20037). Cells were sonicated (Misonix 3000) with a 1/8” microtip probe on ice for alternating intervals of 5 seconds on, 5 seconds off for a total of 2.5 minutes at power level 5.0. The lysed cells were centrifuged at 17,000 x *g* for 45 minutes at 4 °C, and the supernatant was run through a 0.45 µm filter. The protein was applied to a Ni-NTA column (Histrap FF, Cytiva) at 4 °C at a flow rate of 0.8 mL/min on an AKTA start FPLC system (Cytiva). The column was washed with 20 column volumes of 30 mM imidazole in 3X PBS, and the his-tagged protein was eluted with 250 mM imidazole in 3X PBS. The eluate was concentrated with an Amicon 10 kDa centrifugal filter (Millipore UFC8010). The protein was further purified with a weak-anion exchange column (WAX-10 4x250 mm Thermo Scientific) attached to an Ultimate 3000 UHPLC system (Thermo) at 1.0 mL/min in 20 mM tris pH 7.5 with a gradient of 0 to 500 mM NaCl. Fractions of 1 mL were collected and analyzed with SDS-PAGE. The sample was desalted and exchanged into PBS with sephadex G-25 PD-10 column (Cytiva) and concentrated with an Amicon 10 kDa centrifugal filter. Absorbance at 280 nm was converted to concentration in mg/mL using molar extinction coefficients calculated with Expasy ProtParam server [3]. Proteins were stored in PBS with 25% glycerol, flash frozen in methanol/dry ice and stored at -80 °C.

**Mass spectrometry of intact SDC1.** Purified recombinant SDC1 was reconstituted (10-50 μg/ 500 μL) in ammonium formate buffer (10 mM pH 6.8) and dialyzed against 2.5 mM ammonium formate buffer (pH 6.8) at 4 °C using Amicon Ultra 3K MWCO spin dialyzers. The filtrate (50-75 μL) was collected and re-suspended with an equal volume of methanol with 2% formic acid, prior to injection into a LTQ Orbitrap XL (Thermo Fisher).

**MMP-9/Sheddase treatment of SDC1.** TAMRA-labeled SDC1-37 was at a concentration of 10 μM (in 50 mM tris pH 7.4 with 4 mM $CaCl_2$, 150 mM NaCl and 0.05% triton X-100). MMP-9 (R&D Biosystems, 909-MM) was first activated in a 0.1 mg/mL solution of the same buffer by treatment with 1.0 mM 4-Aminophenylmercuric acetate (Sigma, A9563; PAPMA) for 2 h at 37 °C. ADAM-TS1 (R&D Biosystems, 2197-AD) stock was at 0.06 mg/mL. 30 μL of activated MMP-9 or 40 uL of ADAM-TS1 were added to 85 μL of TAMRA-labeled SDC1-37 and incubated at 37 C overnight. For trypsin control, 30 uL of 0.25 % trypsin was added. For the samples treated with MMP-9 and trypsin, His-tagged TAMRA-SDC1 was captured after the reaction with 50 μL of Ni-NTA slurry, washed 3X with 250 μL PBS, and imaged. For ADAM-TS1 treated sample, the TAMRA-SDC1 was reacted while bound to the Ni-NTA resin, washed 3X with 250 μL PBS, and imaged.

**Preparation of 1-bromo per-acetylated xylose**. To a solution of D-(+)-xylose (20.5 g, 136.7 mmol) in acetic anhydride (164.7 mL, 0.83 M) was added sodium acetate (11.2 g, 136.7 mmol) and the resulting reaction mixture was heated to 140 °C. After 5 h, the reaction mixture was allowed to cool to room temperature at which ice-cold water was added. After stirring for 2 h, the reaction mixture was filtered, and the precipitate was purified via recrystallization from water to yield per acetylated xylose 8 (17.8 g, 42%) as a fine white solid. To a cooled (0 °C) solution of the per-acetylated xylose (1 g, 3.14 mmol) in anhydrous $CH_2Cl_2$ (2.66 mL, 1.2 M) was added HBr (1.7 mL, 9.43 mmol) dropwise. The reaction mixture was allowed to stir at 0 °C for 1 h before warming to room temperature. After stirring for 1 h, the reaction mixture was quenched with ice-cold water then diluted with $CH_2Cl_2$. The layers were separated, and the aqueous layer was extracted with $CH_2Cl_2$. The combined organic layers were washed with $NaHCO_3$, brine, dried over anhydrous $Na_2SO_4$, and concentrated *in vacuo*. The crude residue was purified via column chromatography (silica gel, hexanes/EtOAc) to yield 1-bromo per-acetylated xylose (917.1 mg, 86%) as a white solid.

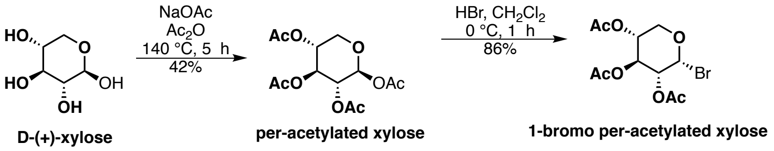


**Preparation of *p*-azidophenol.** Following a previously published procedure, [3], to a cooled (0 °C) solution of 4-aminophenol (1 g, 9.63 mmol) in trifluoroacetic acid (22 mL) was added sodium nitrite (758 mg, 10.99 mmol) slowly. The resulting reaction mixture was allowed to stir for 1 h, at which sodium azide (893 mg, 13.74 mmol) was added over 15 minutes. After 1 h, the reaction mixture was quenched with diethyl ether. Following extraction, the combined organic layers were washed with $NaHCO_3$, brine and filtered through $Na_2SO_4$. The crude residue was purified via column chromatography to afford *p*-azidophenol as a dark brown oil.

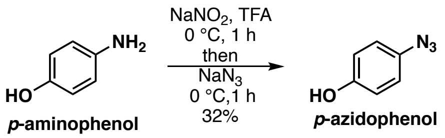


**Preparation of azidoxyloside 1.** To a stirred solution of 1-bromo per-acetylated xylose (319 mg, 2.36 mmol) in 1 M $Na_2CO_3$ (13 mL, 0.18 M) was added TBAB (380.5 mg, 1.18 mmol) followed by a solution of *p*-azidophenol (400 mg, 1.18 mmol) in $CH_2Cl_2$ and the resulting reaction mixture was heated to 40 °C. After 3 h, the reaction mixture was allowed to cool to room temperature then diluted with water. The layers were separated, and the aqueous layer was extracted with $CH_2Cl_2$ (2X). The combined organic layers were washed with $NaHCO_3$, brine, dried over $Na_2SO_4$ and concentrated *in vacuo*. The crude residue was purified via column chromatography to yield the per-acetylated azidoxyloside (288.4 mg, 62%) as a dark brown oil. To a solution of the per-acetylated azidoxyloside (180.8 mg, 0.46 mmol) in methanol (9 mL, 0.05 M) was added potassium carbonate (317.6 mg, 2.3 mmol). The resulting reaction mixture was allowed to stir at room temperature. After 1 h, the reaction mixture was filtered through a pad of silica gel pre-equilibrated with $CH_2Cl_2$. The combined organic layers were concentrated *in vacuo*. The crude residue was purified via column chromatography (silica gel, hexanes/EtOAc/AcOH 10/90/1) to afford the azido xyloside **1** (35 mg, 29%) as an off-white solid.

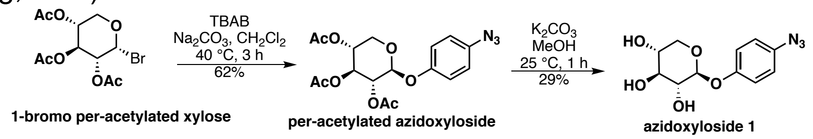


**Preparation of azido-heparin or azido-CS**. We followed a previously published procedure for the synthesis of azide-modified GAGs. [4] A solution of 121.3 mg heparin (Iduron, Hep001) in 1075 µL aqueous buffer (100 mM sodium acetate, 100 mM aniline, pH 5.5) was mixed with 615 µL of 1.05 M solution of 4-(azidomethyl)benzhydrazide in DMSO (123 mg, approx. 80 eqs). The mixture was protected from light and heated in capped Eppendorf tubes on a heatblock at 50 °C for 72 h. The reaction was diluted into 40 mL of PBS, filtered with a 0.45 µm filter (Millex HP) and dialyzed in 10 mM ammonium bicarbonate for 48 h, changing the buffer 3 times. The sample was lyophilized for a yield of 66.8 % (81.0 mg) For synthesis of CS-azide, a solution of 139 mg CS (Sigma, C9819) was prepared in 1230 µL aqueous buffer (100 mM sodium acetate, 100 mM aniline, pH 5.5) and mixed with 705 µL a 1.05 M solution of 4-(azidomethyl)benzhydrazide in DMSO (141 mg, approx. 100 eqs). Reaction conditions and purification were the same as heparin-azide. Yield of CS-azide was 52 % (72.0 mg).

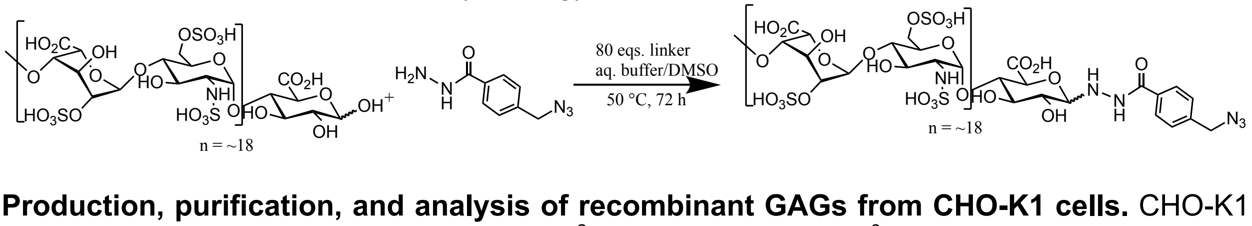


**Production, purification, and analysis of recombinant GAGs from CHO-K1 cells.** CHO-K1 cells were seeded overnight in 10 or 15 $cm^2$ dishes at 10-15 x $10^6$ cells/dish. Compound **1** (30 mM DMSO stock solution) was then added in media at 400 µM for 2-3 days. Following incubation, the conditioned media was collected and xyloside-GAGs were purified by anion exchange chromatography. Media was filtered with a 45 µm filter, then loaded onto a 5 mL Hitrap DEAE sepharose fastflow column (Cytiva, 17515401) equilibrated with PBS at a flow rate of 4 mL/min. The column was washed with 25 column volumes of PBS, then the GAGs were eluted with 6 column volumes of 1.5 M NaCl in PBS. The eluate was dialyzed for 24 hr against 10 mM ammonium bicarbonate in 3500 MWCO tubing (Snakeskin, ThermoFisher, 68035), changing the buffer 3 times. Following dialysis, the sample was lyophilized and reconstituted in DPBS containing calcium and magnesium (Corning, 21-030-CM), with 50-100 μL of DPBS per every 10 mL of media purified. To isolate heparan sulfate (HS) and chondroitin sulfate (CS) components of the GAGs, the purified GAGs were split and digested with heparinase or chondroitinase enzymes. To purify HS GAGs, CS was digested using 0.5 U/mL chondroitinase ABC (Sigma, C3667) in pH 7.5 DPBS (Corning, 21-030-CM) supplemented with 50 mM sodium acetate. To purify CS GAGs, HS was digested with a mixture of heparinase I, II, and III at 2.5 U/mL each (heparinase II, Sigma H6512; heparinase I and III, Sigma H3917). Disaccharides resulting from the enzymatic treatment were removed by four repeated rounds of buffer exchange at 16,000 x *g* for 20 minutes on an Amicon 3 kDa MWCO centrifugal filter (Millipore, UFC500). Filtrate was harvested for subsequent disaccharide analysis. Potential remaining partially digested oligosaccharides were removed with a Sephadex G-25 PD-10 column (Cytiva).

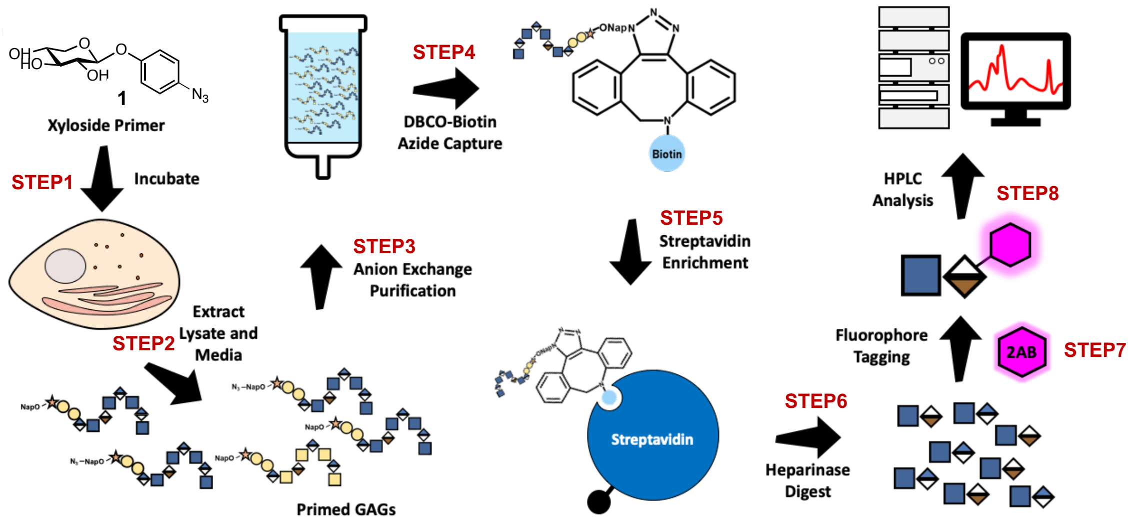


**Scheme S1. Workflow for the production and analysis of recombinant GAGs.** The azidoxyloside primer **1** was incubated with CHO-K1 (wt) cells for 2-3 days at 37°C. The conditioned media and the cell extracts were collected, purified by anion exchange chromatography (manual columns with DEAE resins), and the eluate from the high-salt elution was desalted (PD-10 columns). The harvested GAGs were then reacted with dibenzocyclooctyne (DBCO)-biotin in solution and azide-reacted GAGs were immobilized on streptavidin beads. GAGs were then harvested with a cocktail of heparinases or chondroitinases on-beads, and the resulting solution was reacted with 2-AB fluorophore and analyzed by HPLC.

**Preparation of chondroitinase ABC and heparinase cocktail.** A mixture of heparinases was prepared by adding 1 mL PBS (with Ca and Mg) per 100 Unit bottle of Heparinase I and III (Sigma, H3917). After mixing, 1 mL of the above solution was added to 100 units of heparinase II (Sigma, H6512). Chondroitinase ABC (Sigma, C2905) stock solutions were prepared at 5.0 U/mL in 60 mM tris, 50 mM sodium acetate, pH 7.9 buffer. Aliquots of the enzymes were stored at -80 °C.

**Enzymatic digestion of GAGs.** GAGs (soluble or covalently attached to beads) were resuspended in 400 μL of PBS (+ Ca, Mg). Then, 10 μL of the heparinase cocktail was added and mixed thoroughly. The samples were rotated at 37 °C for 18 h, and then centrifuged (2000 rpm, 2 min). The supernatant was then transferred to Amicon 3K MWCO spin filters (14,000 x g, 30 mins) to separate undigested material. The digested samples (eluate) were lyophilized and stored at -80 °C.

**GAG disaccharide labeling.** Disaccharides were fluorescently labeled with a 1.0 M solution of $NaBH_3CN$ and 0.2 M anthranilamide (in 30% glacial acetic acid/DMSO) which was prepared immediately before use. To each dried digested sample was added 35 µL of the labeling mixture, which was centrifuged at 6,000 x g for 3 minutes and transferred to fresh Eppendorf tubes. The samples were heated at 70 °C for 2 hours. A column for each sample was equipped with Whatman Grade 3 filter paper pre-equlibrated with water (1 mL), 30% acetic acid (2.5 mL), and acetonitrile (2 mL). Sample was added to the filter paper and allowed to adsorb for 15-20 minutes at room temperature, then washed with 2 mL of acetonitrile. The acetonitrile wash was repeated four times. Each column was placed in 15 mL collection tube. Each column was pulse centrifuged ~10 s before being transferred to newly labeled 15 mL conical tube. Each digested sample was eluted by adding 500 µL of water. Then an additional 500 µL of water was added and the sample centrifuged for ~10 s. The samples were stored at -20°C.

**GAG disaccharide analysis** Fluorophore labeled samples were diluted into water and injected onto a Propac-PA1 column (4 x 150 mm, Thermo) attached to a Propac-PA1 guard column (4 x 50 mm, Thermo) on an Ultimate 3000 UHPLC system (Thermo) at a flow rate of 1.0 mL/min, with a gradient from 0 to 1.0 mM NaCl in 50 mM NaOH. Fluorescent excitation and emission wavelengths were 348 and 440 nm, respectively. Retention times for HS disaccharides were confirmed using a heparin disaccharide standard mix (Galen, HD Mix).

**CuAAC click reactions.** Reactions took place in PBS buffer at pH 6 to 8, with concentration of the alkyne-containing protein at approximately 25 µM and the GAG-azide (or TAMRA-azide) at 900 µM. Some reactions were ran with the core protein in excess (30 µM) and GAG-azide at approximately 10 µM. Aminoguanidine hydrochloride was added to a concentration of 5.0 mM [5]. Copper sulfate final concentration was 320 µM, Tris(3-hydroxypropyltriazolylmethyl)amine (THPTA) was 1600 µM, and sodium ascorbate was 21 mM. Alkyne and azide reactants were mixed, along with aminoguanidine, and the mixture of click reagents (copper sulfate, THPTA, and sodium ascorbate) were added last. To prepare the click reagents, we first prepared a mixture of 1.2 mM copper sulfate and 6 mM THPTA in water. This was sparged with nitrogen for 5 minutes. A 200 mM solution of sodium ascorbate was prepared from water that had likewise been sparged with nitrogen. The sodium ascorbate solution was vortexed with the pre-complexed mixture of copper sulfate and THPTA and diluted into the reaction. The reaction was shaken in a sealed Eppendorf tube at 300 rpm at 37 ˚C for 2 hours. Protein-GAG conjugate was purified on a weak anion exchange DEAE column (WAX-10 4x250 mm Thermo Scientific) on an Ultimate 3000 UHPLC system at 1.0 mL/min in 20 mM tris pH 7.5 buffer. A sodium chloride gradient from 0 to 400 mM NaCl was applied over 15 minutes, at which point the NaCl concentration was raised to 1.75 M. The reaction product was collected in the high-salt eluting fractions. To remove excess GAG, the collected DEAE fractions were loaded onto a Ni-NTA column (Histrap FF, Cytiva) and washed with 20 column volumes of 1 M NaCl with 20 mM Tris at pH 7.5. The GAG-linked protein was eluted with 5 column volumes of 400 mM imidazole, 500 mM NaCl, in 25 mM Tris at pH 7.5. This was exchanged into PBS with a sephadex G-25 PD-10 column (Cytiva), and concentrated to approximately 50 µM with an Amicon 10 kDa centrifugal filter. Concentration was measured with a nanodrop apparatus and absorbance at 280 nm was converted to molar concentration [3]. For CuAAC conjugation to TAMRA azide 5-isomer (Lumiprobe, 47130), the procedure was the same, with 10 mM TAMRA azide 5-isomer in DMSO diluted into the reaction mixture to give 200 µM final concentration. After 1 to 2 hours an aliquot of the reaction was mixed with SDS-PAGE sample buffer containing 10% 2-mercaptoethanol and loaded onto the gel. GAG-azides were labeled with alexafluor-488-yne (Sigma, 761621) using CuAAC in a similar manner as described above, with GAGs extracted from a 15 $cm^2$ dish dissolved in 200 uL PBS and reacted with 200 µM alexafluor-488-yne. Excess fluorophore was removed with a PD minitrap G-10 column (Cytiva). Copper-free click chemistry labeling of GAG-azides with DBCO-PEG4-fluor 545 (Sigma, 760773) was performed by incubating GAGs with 200 µM DBCO-PEG4-fluor 545 for 20 h at 37 ˚C.

**HPLC monitoring of CuAAC reactions and SEC GAG analysis** Click reactions between alkyne-displaying proteins and azide-GAGs were analyzed with a Propac WAX-10 column (4 x 250 mm on an Ultimate 3000 UHPLC system (Thermo) at a flow rate of 1.0 mL/min with a gradient ranging from 0-1.75 M NaCl (20 mM tris pH 7.5) with fluorescent detection ($\lambda_{ex}$=280 nm, $\lambda_{em}$=350 nm). Analytical size exclusion chromatography was performed with a MabPac SEC-1 column (Thermo, 5 x 300 mm) at a flow rate of 0.25 mL/min in PBS at pH 7.15, with an injection volume of 10 $\mu$L, with fluorescence detection settings adjusted for the respective fluorophore of interest.

**Biotinylation of FGF2**. A HEPES buffered (200 mM, pH 8.4, 50 $\mu$L) solution of FGF2 (100-150 $\mu$g, 35 $\mu$L) was prepared with heparin (20 mg/mL in $H_2O$, 10 $\mu$L) and sulfo-NHS-biotin (4 mg/mL $H_2O$, 5 $\mu$L) was reacted for 2 hrs (final volume: 100 $\mu$L) at room temperature. A glycine solution (20 $\mu$L, 10 mg/mL) was then added to stop the reaction. The solution was then purified and eluted (3 M NaCl, 0.2% BSA, 20 mM HEPES, pH 7.4) using a heparin-sepharose column (GE Healthcare 17-0998-01) and stored at 4°C prior to use.

### BIOLOGICAL METHODS

**Flow cytometry.** Cells were harvested using non-enzymatic cell dissociation buffer (MDA-MB-231) or EDTA-free 0.05% trypsin (remodeled mESCs) into 96-well round bottom plates. After centrifugation at 500xg for 5mins at 4°C, the supernatant was flicked off and cells were fixed in 4% PFA/PBS for 10 mins at RT. Cells were washed in 1X DPBS twice before incubation with antibodies in 3% BSA for 1h, on ice. If secondary antibodies were being used, cells were washed twice in 1X DPBS before secondary antibody incubation for 1h, on ice, in dark. Cells were washed twice with 1X DPBS before re-suspension in 5% FBS in PBS and ran on an BD Accuri C6. Data analysis was performed using FlowJo and Prism 9.

**Neuronal differentiation of mouse embryonic stem cells (mESCs), extended**. To form embryoid bodies (EBs), a cell suspension of $3x10^5$ cells/ml was prepared in mESC maintenance media and added in 20ul droplets onto the top of a cell culture dish filled with sterile PBS and grown for 3 days as hanging drops. On day 3, EBs were washed from the lid and split between a 6-well plate and incubated in N2B27 media ± soluble heparin (T0). Media was changed every two days and EBs were kept in suspension. On day 9 (T6 in N2B27), EBs were harvested and allowed to settle before media was aspirated and cells were fixed using 4% PFA/PBS for 10 mins at RT. For immunofluorescence, fixed cells were distributed into 8-well chamber slides (*Nunc™ Lab-Tek™ II Chamber Slide™*) for immunofluorescence staining.

**Immunofluorescence staining.** After fixation, cells were blocked in ICC buffer (3% BSA, 0.1% Triton in PBS) for 1h at RT with rocking. Cells were then incubated with primary antibodies (See Table 2) in ICC buffer for 1h at RT, or overnight at 4°C, with rocking. Cells were washed twice with 1X DPBS before incubation with secondary antibody (1:1000, see Table 2) in ICC buffer, for 1h at RT with rocking in the dark. Nuclei were stained using Hoescht 33342 for 5 mins at RT with rocking. Cells were washed twice with PBS before imaging. EBs were removed from 8-well chamber slides at this point and mounted on microscope slides in Antifade mounting solution (ThermoFisher). Slides were allowed to dry overnight at 4°C before imaging. All cell imaging was performed on a benchtop EVOS M5000 fluorescence microscope (Thermo Fisher Scientific).

**FGF2 stimulation assay.** Cells were seeded at $1x10^6$ cells per well in a 0.1% gelatin-coated 6-well plate and allowed to adhere overnight. The following day, cells were serum starved with serum free mESC maintenance media overnight. Remodeled $EXT1^{-/-}$ were incubated with 10μM cholPEGNTA for 1 hr at 37°C before on-cell complexation with 2μM SDC1 constructs (1h, 37°C). Soluble PGs (2 μM) were pre-incubated for 1h before excess was washed away. Stimulation was performed by adding 25ng/mL recombinant human basic FGF (Cell Signalling, #8910) in KO-DMEM to cells for 15 mins at 37°C. Cells were scraped before lysis in ice cold 1X RIPA (Cell Biolabs, #AKR-191) supplemented with protease inhibitor cocktail and 1mM PMSF. Protein concentration was quantified using the DC Protein Assay (BioRad, #5000111) and normalised in PBS. 10ug of lysate was mixed with SDS-PAGE sample buffer containing 10% 2-mercaptoethanol and loaded onto the gel. Membranes were first probed for phosphor-ERK1/2 and α-tubulin, stripped with Restore™ PLUS Western Blot Stripping Buffer (ThermoFisher, #46430) for 10 mins and re-probed for ERK1/2 and α-tubulin.

**Knockdown of SDC1 via RNAi.** The TriFECTA RNAi kit (IDT Technologies design ID hs.Ri.SDc1.13) encoding three dicer-substrate short interfering RNAs (DsiRNAs) against human SDC1 was obtained. MDA-MB-231 cells were treated with a pooled mixture of all three DsiRNAs (100 or 200 nM). Parameters for transfection were optimized using the fluorescent non-targeting control and transduction control, and performed using Lipofectamine RNAiMAX (Invitrogen) and incubated at RT for 5 minutes before DsiRNA-lipid complex was added drop-wise to cells. A scrambled DsiRNA that does not recognize any human sequence was used as a negative control, alongside a fluorescently labelled TYE563 transfection control. 24-well plates were coated in 1X poly-D-lysine (15 mins, RT), washed twice with 1X DPBS before incubation with 10μg/ml vitronectin (VN) overnight at 4°C, rocking. At 48h, the plate was washed twice with PBS, followed by blocking for 1h at 37°C in DMEM supplemented with 1% BSA. At 48h, MDA-MB-231 cells harvested using non-enzymatic cell dissociation buffer (PeproTech #CPD-125) at a density of $5x10^4$ cells per well onto VN matrices. Cells were allowed to adhere for 2h at 37°C before being washed with 1X PBS and fixed in 4% PFA/PBS. Cell spreading was visualized using rhodamine-conjugated phalloidin (1:40) in ICC buffer, following to immunofluorescence staining protocol.

| DsiRNA | Cross-reacting transcript | Location | Exon |
|---|---|---|---|
| 1 | NM_002997 | 3' UTR | 5 |
| | NM_001006946 | 3' UTR | 6 |
| 2 | NM_001006946 | CDS | 4 |
| | NM_002997 | CDS | 3 |
| 3 | NM_002997 | CDS | 3 |
| | NM_001006946 | CDS | 4 |

**Generation of $SDC1^{KO}$ cells.** MD-MBA-231 cells were seeded at a density of $2.5x10^5$ cells per well in a 6-well plate and allowed to adhere overnight (37°C, 5% $CO_2$, ~18 hr) before spinoculation (2000 rpm, 2 hr, 37°C) with guideRNA (gRNA) targeting SDC1 exon 5 (gRNA sequence GTTCCGGCGGTCAGGCTCCA) ordered from the GenCRISPR™ Plasmid Collection. Cells were kept in lentiviral-supplemented DMEM overnight before being changed to DMEM/10% FBS and allowed to expand for 4 days. Transduced cells were selected with 5 μg/mL puromycin.

**Quantification of cell spreading.** Cells were imaged on an EVOS M5000 fluorescence imager (Thermo Fisher) and analyzed using ImageJ. Individual channels were converted to 8-bit greyscale and segmented with the threshold function to generate a black and white image of spreading of cell cytosol. The ‘analyze particles’ function was then used (size 150-infinity) to quantify cell spreading. This was performed on both DAPI and rhodamine-conjugated phalloidin images. The cytosolic area from phalloidin stained images was divided by DAPI count per image to quantify extent of spreading. Cell spreading was normalized to non-treated cells to generate a percentage of cell spreading. Graphing and statistical analyses were performed using Prism 9.

### ADDITIONAL DATA

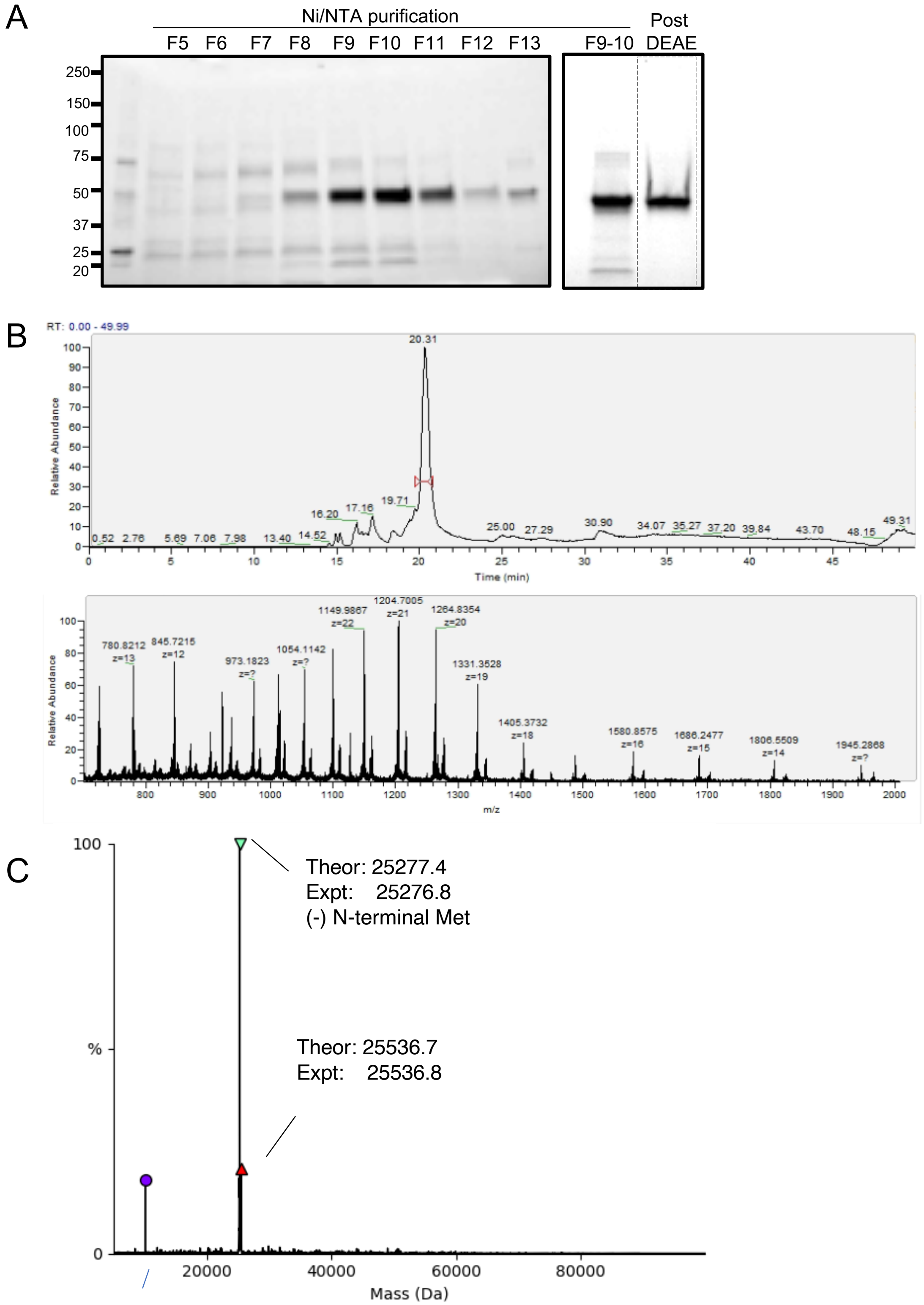


**Figure S1. Characterization of SDC1$_{37}$.** (**A**) SDS-PAGE analysis of the protein following Ni/NTA-based purification and DEAE purification. (**B,C**) Intact mass analysis shows the expected molecular weight of the construct.

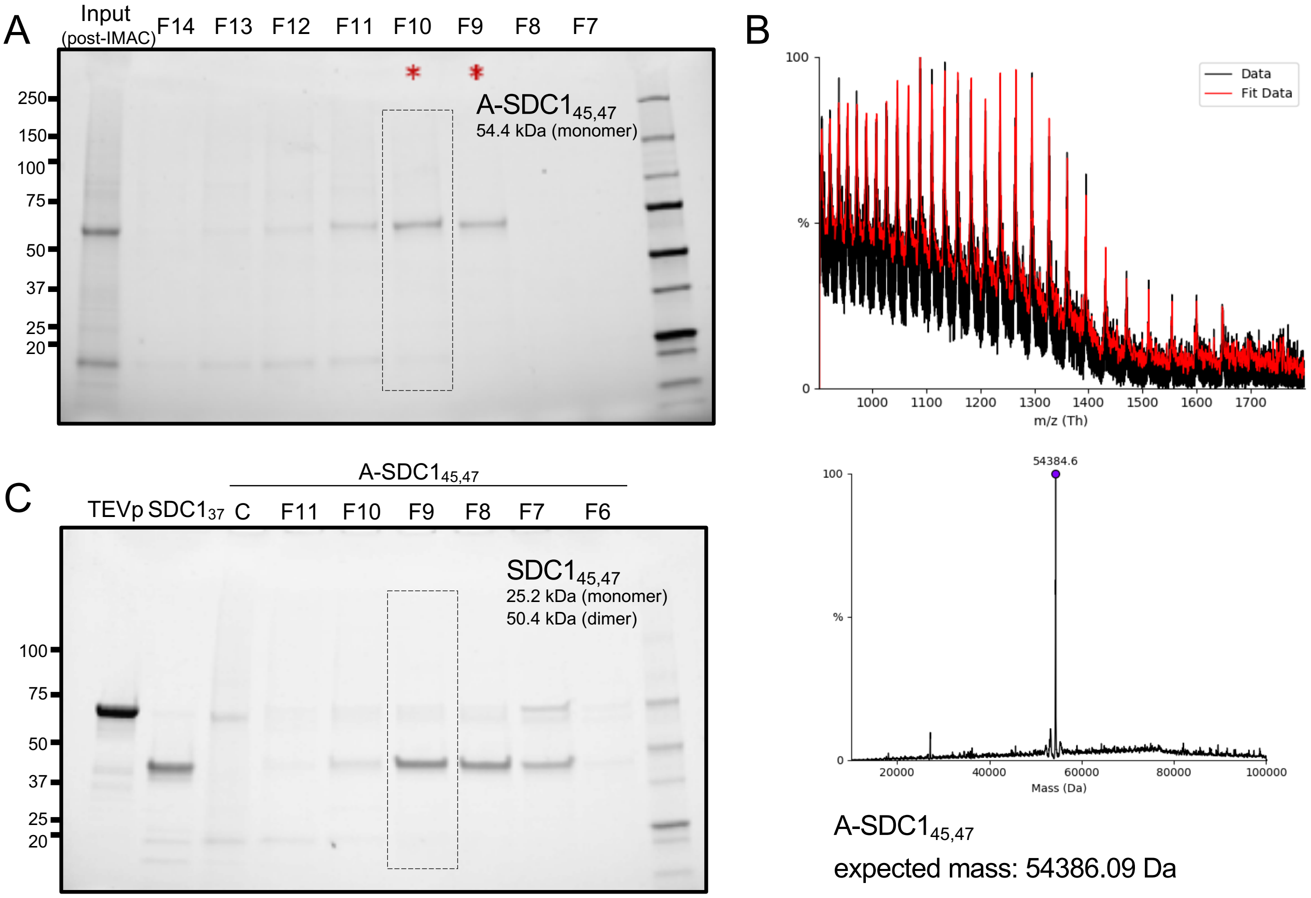


**Figure S2. Characterization of A-SDC1$_{45,47}$.** (**A**) SDS-PAGE analysis of the protein following Ni/NTA-based purification. (**B**) Intact mass analysis shows the expected molecular weight of the construct. (**C**) SDS-PAGE analysis of TEV-mediated cleavage and subsequent purification of A-SDC1$_{45,47}$ into the divalent SDC1$_{45,47}$.

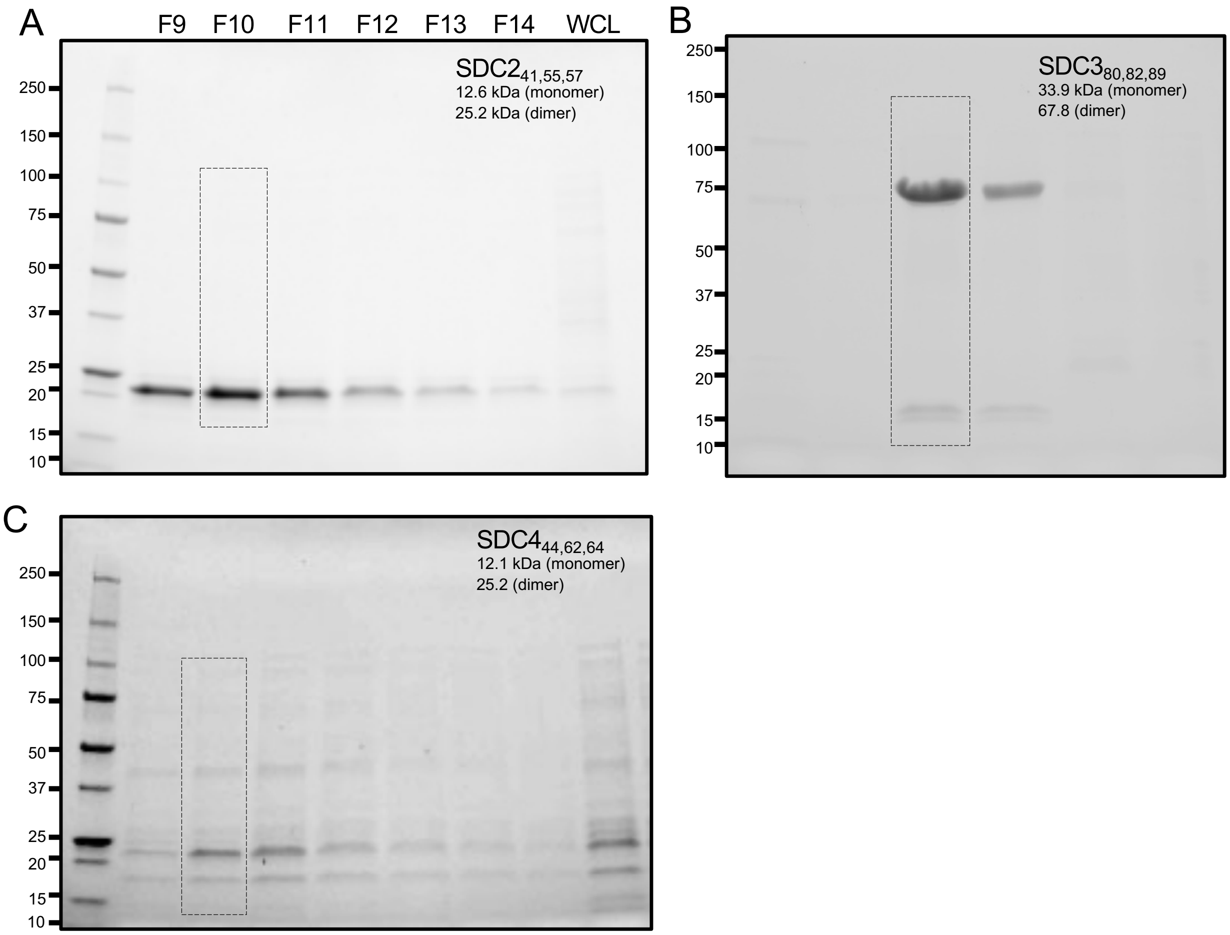


**Figure S3. SDS-PAGE analysis of fractions from Ni/NTA purification for** (**A**) $SDC2_{41,55,57}$. (**B**) $SDC3_{80,82,89}$ (**C**) $SDC4_{44,62,64}$. All three syndecans were expressed as deglycosylated proteins that migrate as dimers as previously reported [6].

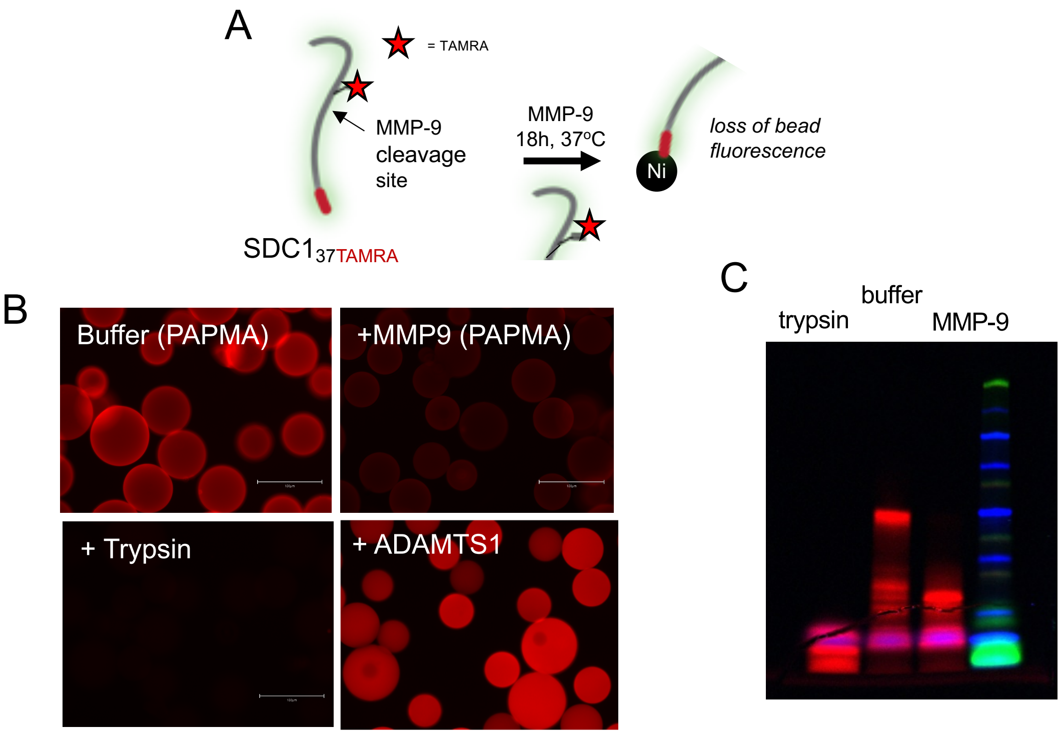


**Figure S4. Sheddase activities of engineered SDC1$_{37TAMRA}$.** Engineered SDC1 retains ability to be selectively cleaved by known proteases. **(A)** Schematic of on-bead cleavage experiment. SDC1$_{37TAMRA}$ was immobilized on Ni/NTA beads and treated with MMP-9, ADAMTS1, or trypsin (positive control) proteases. Loss of on-bead fluorescence is deemed as a positive cleavage event. (**B**) Fluorescence microscopy images of beads following treatment with proteases or buffer with *p*-aminophenylmercuric acetate (PAPMA, required to activate MMP) shows SDC1$_{37TAMRA}$ is only cleaved by MMP9 and trypsin. (**C**) Fluorescence SDS-PAGE analysis of the fragments released from the beads.

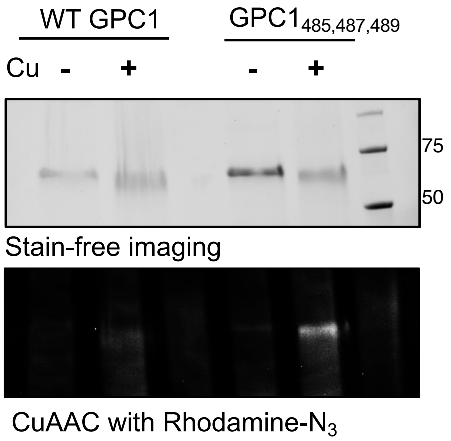


**Figure S5**. **Expression of GPC1$_{485,487,489}$.** Wild-type (wt) and propargyl lysine-incorporating mouse glypican-1 (GPC1) ectodomain (244-529) at residues 485, 587, and 489 were expressed in BL21 *E. coli* cells using EcoRI/HindIII restriction sites and subjected to CuAAC with Rhodamine-azide.

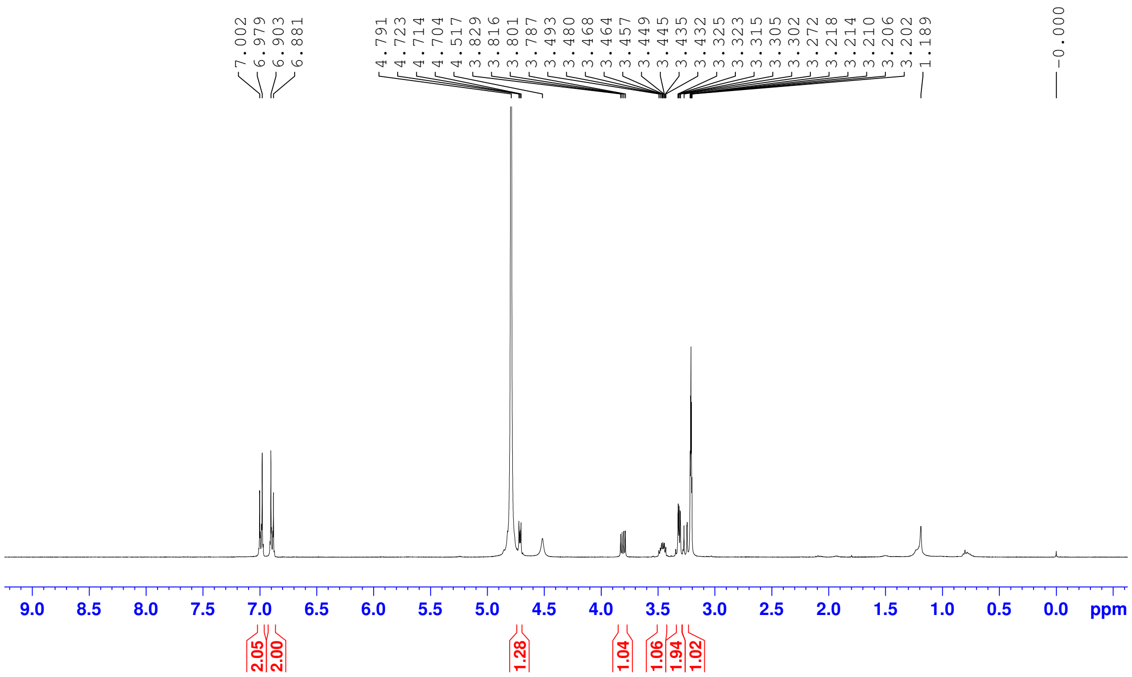


**Figure S6. [1]H NMR analysis of azidoxyloside 1 (400 MHz, $CD_3OD$).**

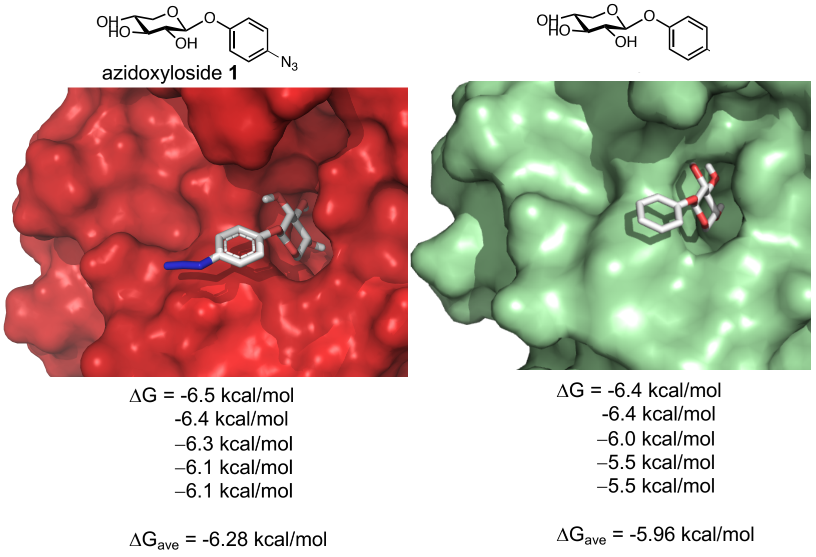


**Figure S7. Molecular docking simulations of compound 1 (left) or its non-azide derivative (right) into the active site of β4GalT7.** Compounds were individually docked into the active site region of β4GalT7 in complex with the UDP-galactose donor (PDB ID 4M4K). [7] Top poses for the docked xylosides showed positioning of the xylose C4 oxygen within 3-4 Å of the C1 atom on the galactose ring of UDP-galactose. The binding affinity of the five lowest energy poses for each simulation was averaged. Comparison of the average binding energies showed the addition of azido group into the aromatic ring component did not exact a substantial binding penalty.

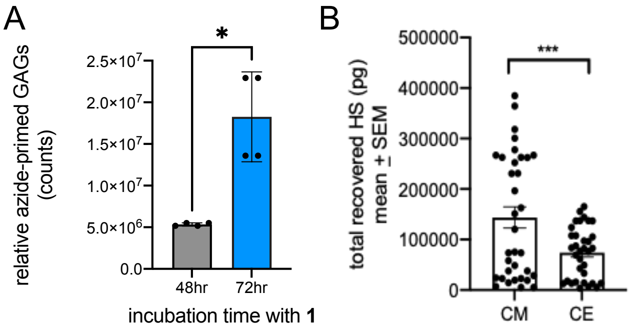


**Figure S8**. **Characterization of recombinant GAGs isolated from CHO-K1 cells**. (**A**) Increased incubation periods lead to larger amounts of azide-primed GAGs. (**B**) Cellular HS isolated from CHO-K1 wild-type cells is mostly present in conditioned media (CM) compared to cellular extracts (CE).

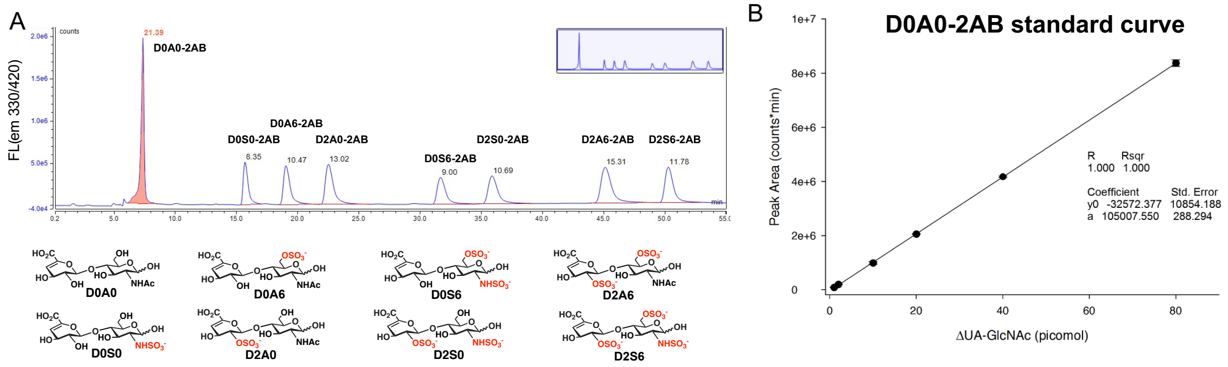


**Figure S9. Analysis of HS disaccharide composition and normalization to absolute amounts.** (**A**) HPLC chromatogram of 2AB-functionalized HS disaccharide standards composed of the eight most abundant sulfation patterns (top). Nomenclature for disaccharides derived from Lawrence et al. [8] (**B**) A standard curve of fluorescence peak area vs. picomols of D0A0 (ΔUA-GlcNAc) was constructed in order to aid in the determination of mols of disaccharide produced.

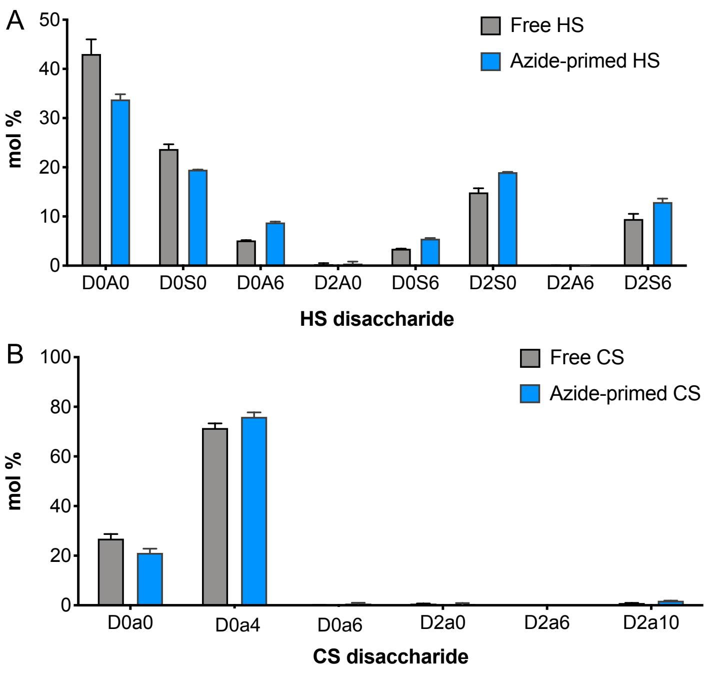


**Figure S10. Recombinant azide-primed GAGs recapitulate sulfation patterns of host GAGs.** (**A**) HS and (**B**) CS disaccharide composition of free versus azide-primed HS and CS, respectively, show similar patterns.

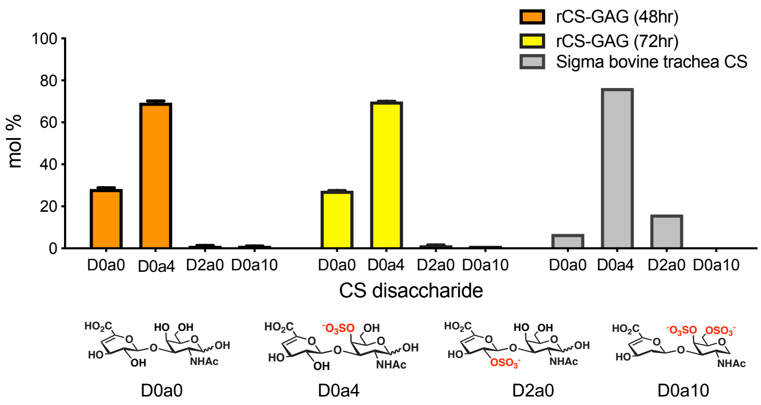


**Figure S11. Recombinant azide-primed CS are similar despite differing incubation times.**

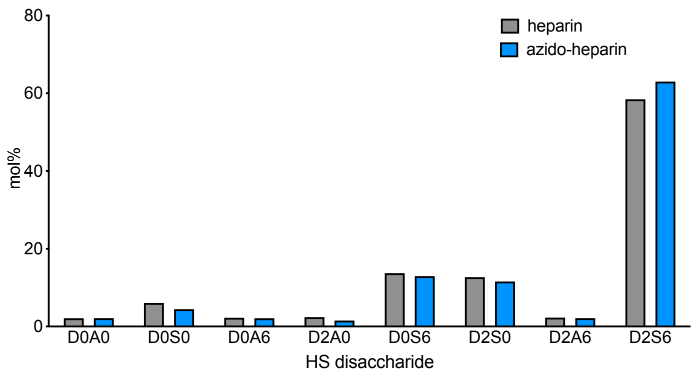


**Figure S12. Azido-heparin mimics the composition of native heparin.**

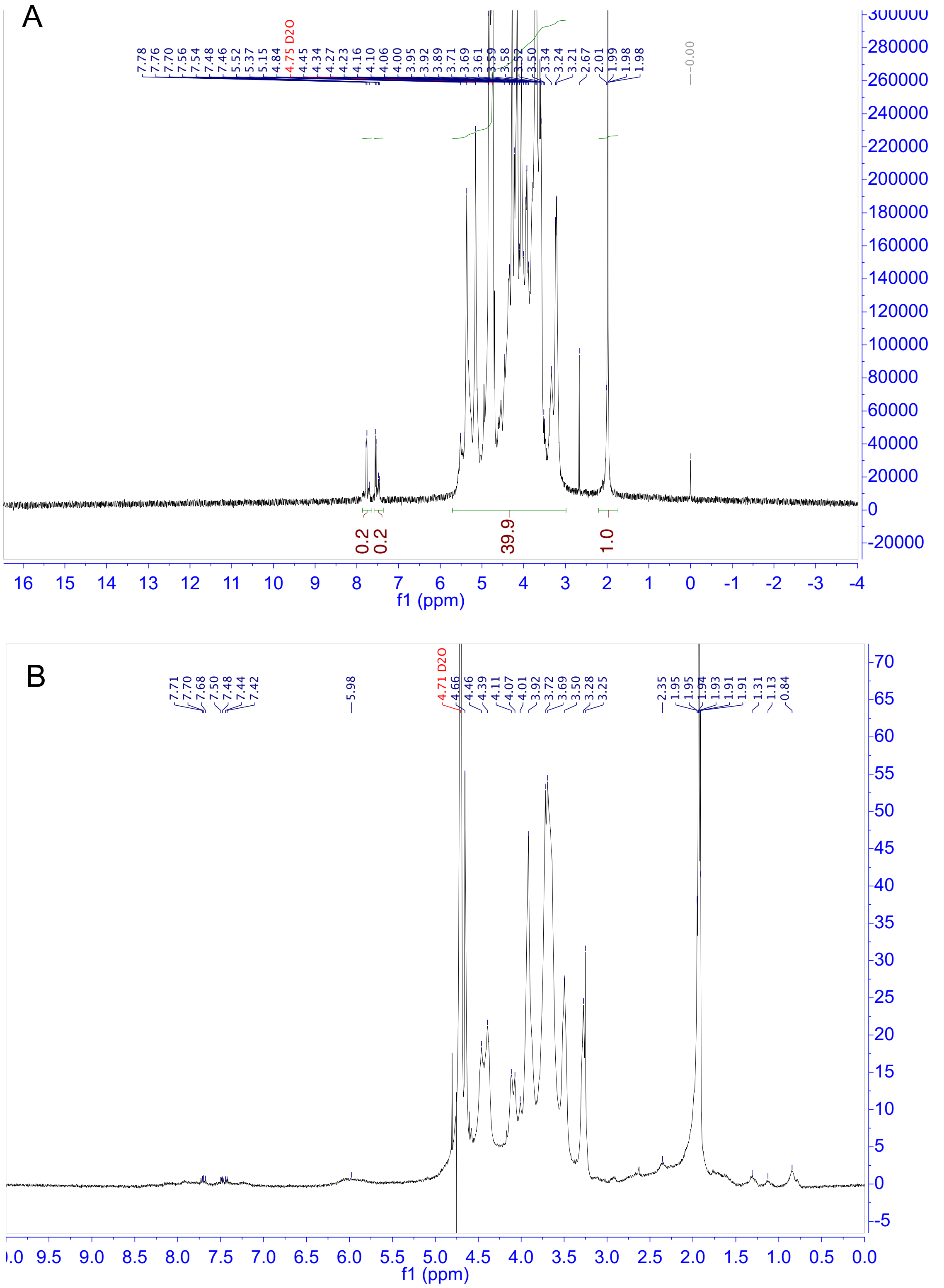


**Figure S13.** **¹H NMR (400 MHz, $D_2O$) analysis of azido-heparin (A) and azido-CS (B).**

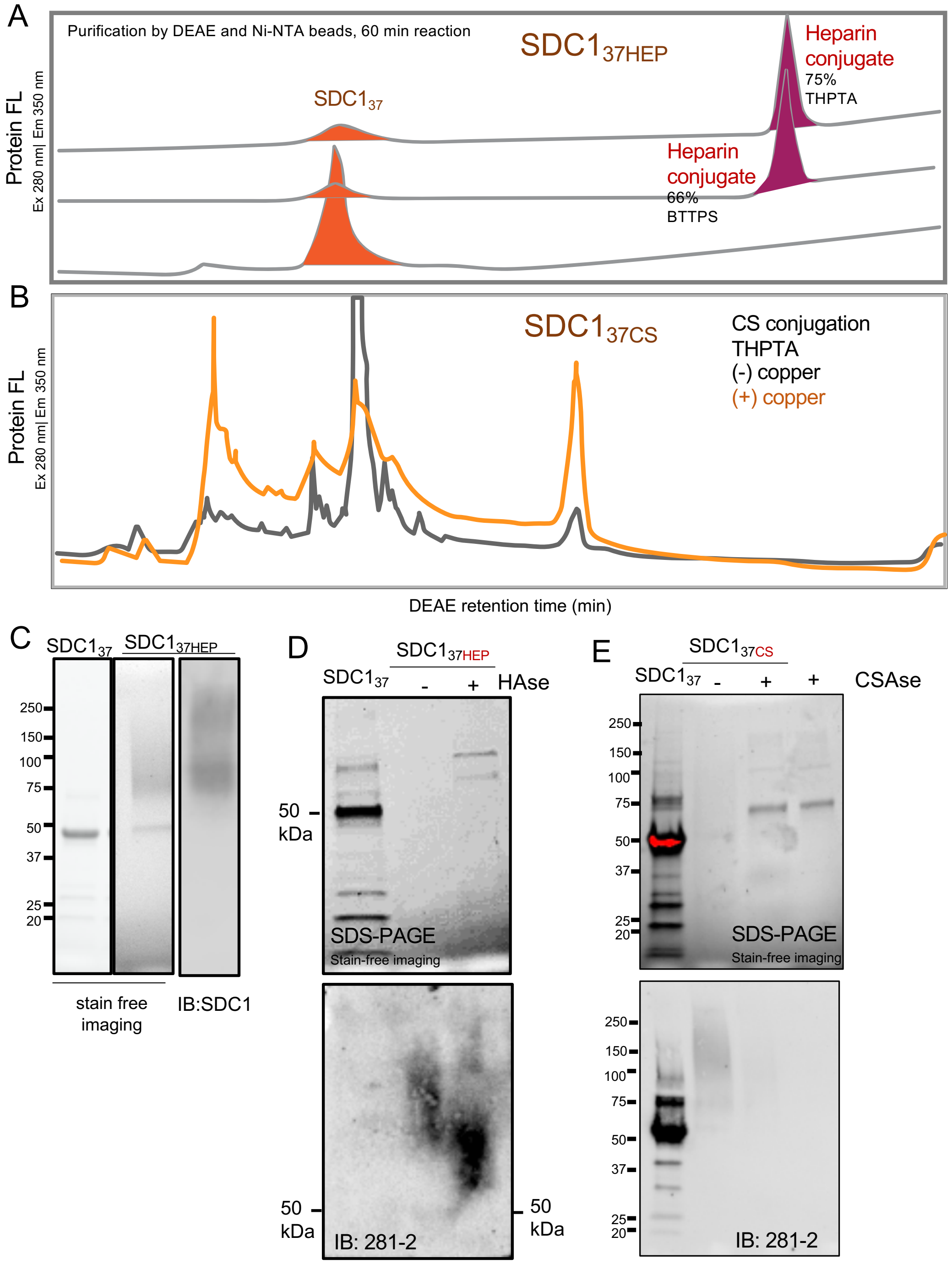


**Figure S14. Additional data for SDC1$_{37}$ conjugated to either azido-heparin (A, C, D) or azido-CS (B, E). (A)** DEAE chromatography of crude reaction mixtures shows greater conversion to SDC31HEP using THPTA as a catalyst for CuAAC over BTTPS. (**B**) THPTA-catalyzed conjugation to form SDC137CS. (**C**) Stain-free imaging of SDC1$_{37HEP}$ conjugates shows a diffuse, higher molecular weight (75-150 kDa) and weak signals compared to localized and strong band observed for SDC137. Western blotting using anti-SDC1 (281-2) confirms the presence of the conjugate. (**D**) Treatment of SDC1$_{37HEP}$ with heparinase shows digestion into lower MW band close to starting material (50 kDa). (**E**) Chondroitinase (CSAse) treatment of SDC1$_{37CS}$ significantly digests material to a lower MW band.

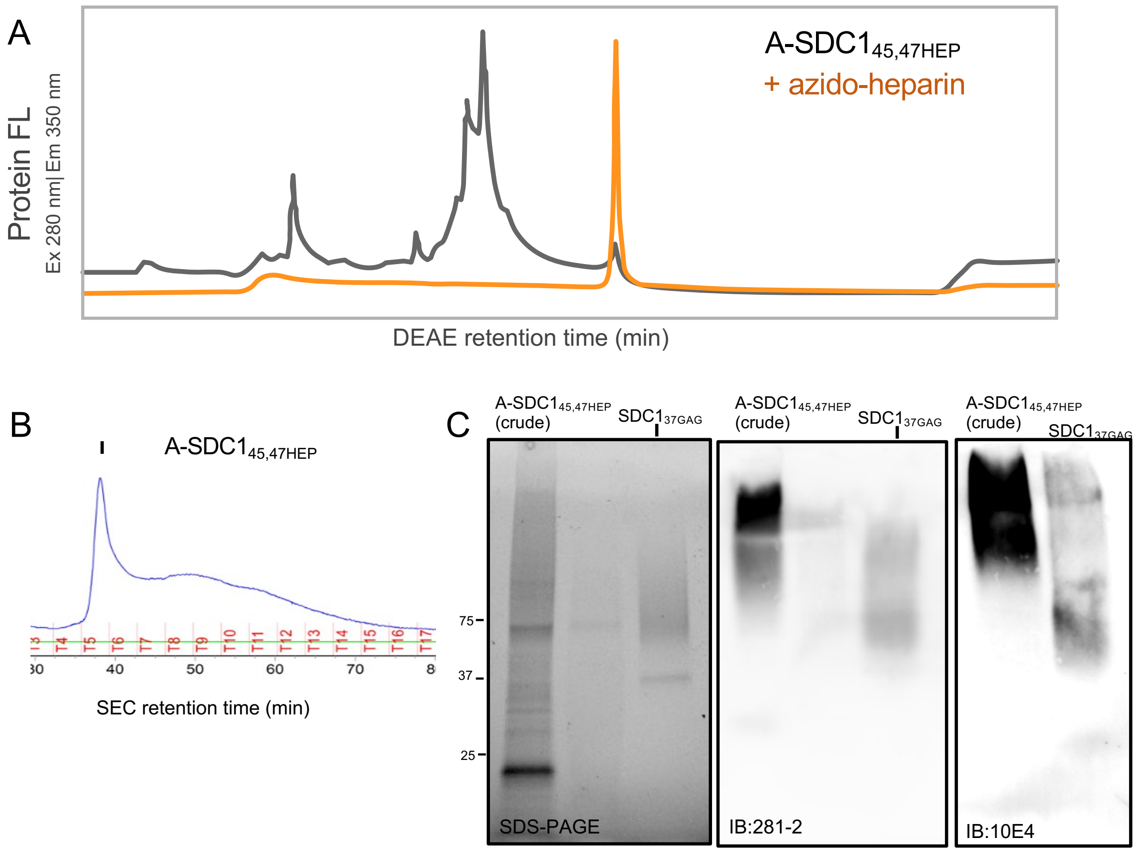


**Figure S15. A-SDC1$_{45,47}$ conjugation to azido-GAGs. (A)** Anion exchange chromatography of the core protein before (black) or after (orange) CuAAC conjugation to azido-heparin shows quantitative conversion to a more anionic molecule. (**B**) Size exclusion chromatography of the protein following anion exchange chromatography and (**C**) subsequent analysis by SDS-PAGE (left panel) and Western blotting (middle, right panels) shows that the divalent conjugate A-SDC1$_{45,47HEP}$ migrates much higher compared to the monovalent SDC1$_{37HEP}$. Note: GAG-conjugated PGs are not very visible by stain-free imaging on SDS-PAGE gels, but they are readily observed by Western blotting when probed using an anti-SDC1 antibody (clone 281-2) or an ant-HS antibody (clone 10E4).

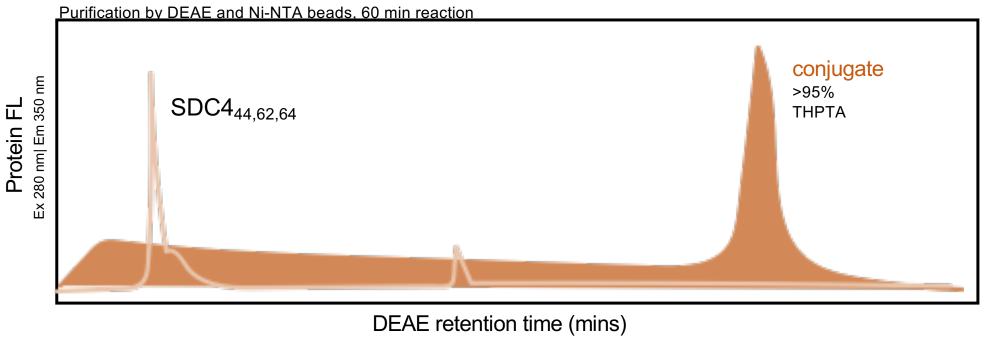


**Figure S16. SDC4$_{44,462,64}$ conjugation with azido-heparin.** Anion exchange chromatography trace of the core protein (light orange trace) and the crude reaction (filled orange) following CuAAC with azido-heparin. The conjugate product elutes with high salt concentrations. There is >95% conversion of the starting material to the conjugate.

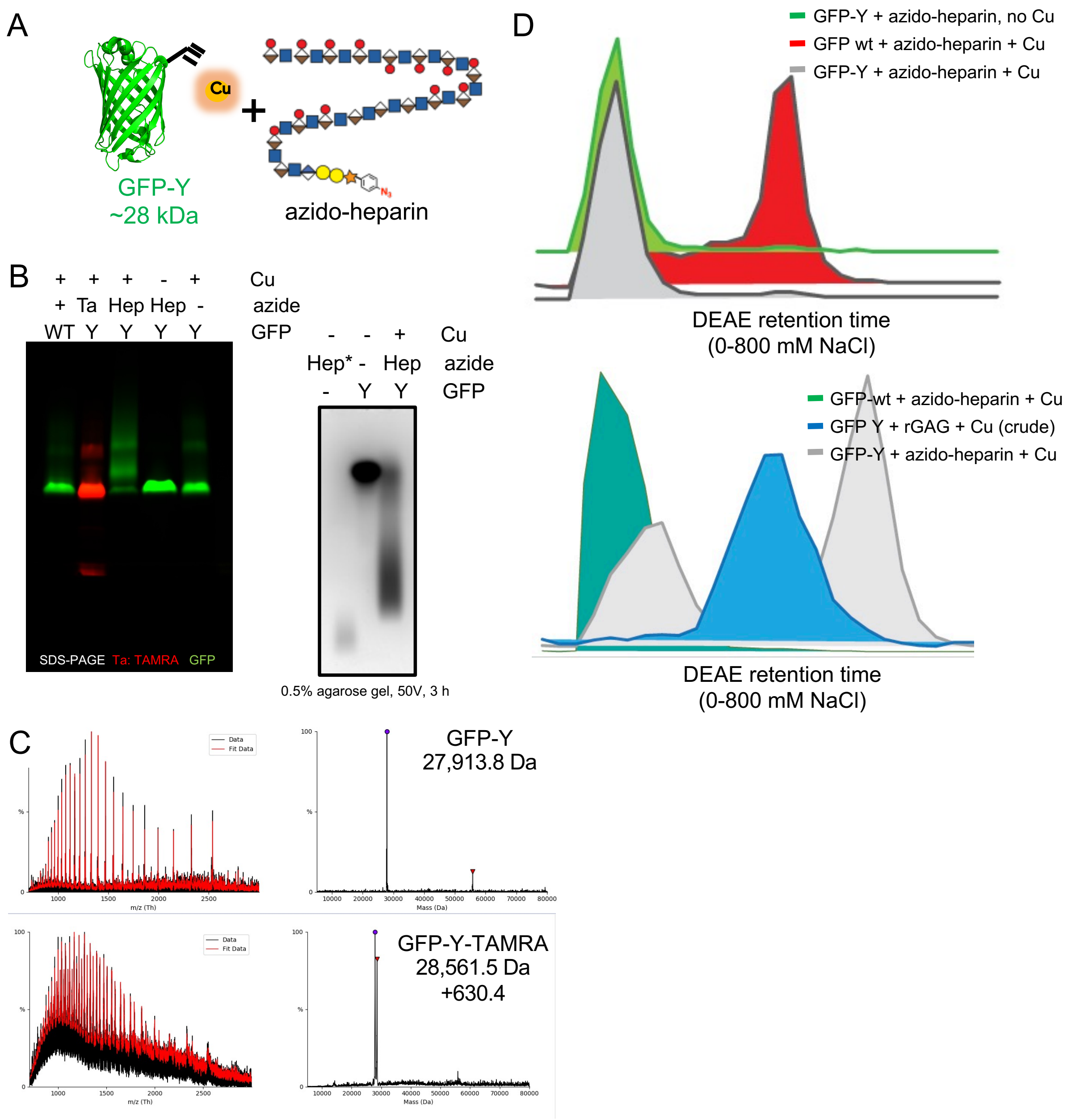


**Figure S17. Conjugation of pPY-containing GFP-Y to azidoGAGs. (A)** Cartoon representation of the reaction. (**B**) Fluorescence visualization of SDS-PAGE gel (left) or agarose gel (right) following CuAAC of WT GFP or GFP-Y with TAMRA-azide fluorophore (Ta) or azide-heparin (Hep). Both types of gel analyses result in diffuse higher molecular-weight bands upon reaction with azide-heparin. In contrast, reaction with a small molecule TAMRA-azide results in a localized band. (**C**) Intact mass spectrometry analysis of GFP-Y and resulting conjugate with TAMRA-azide. (**D**) Fluorescence-based anion exchange chromatography analysis of crude reactions show the formation of more anionic conjugate products with azido-heparin (top, red) or azide-primed recombinant GAGs (bottom, blue).

**Table S2. ELISA data for $\alpha_v\beta_3$ integrin binding to SDC1 ectodomains.**

| | $SDC1_{37}$ | $SDC1_{37HEP}$ | $SDC1_{37rHS}$ | $SDC1_{45,47}$ | $SDC1_{45,47HEP}$ | $SDC1_{45,47rHS}$ | $SDC1_{37CS}$ |
|---|---|---|---|---|---|---|---|
| log(agonist) vs. normalized response -- Variable slope | | | | | | | |
| Best-fit values | | | | | | | |
| LogEC50 | -0.7455 | -1.876 | -1.440 | -0.8562 | -2.139 | -1.484 | -1.412 |
| HillSlope | 1.437 | 1.544 | 1.098 | 1.585 | 1.221 | 1.058 | 1.811 |
| **$EC_{50}$** | 0.1797 | 0.01329 | 0.03635 | 0.1393 | 0.007257 | 0.03280 | 0.03874 |
| 95% CI (profile likelihood) | | | | | | | |
| LogEC50 | -0.7922 to -0.6989 | -1.896 to -1.857 | -1.490 to -1.389 | -0.8867 to -0.8255 | -2.199 to -2.079 | -1.532 to -1.436 | -1.672 to -1.141 |
| HillSlope | 1.269 to 1.633 | 1.460 to 1.635 | 0.9910 to 1.223 | 1.438 to 1.756 | 1.056 to 1.428 | 0.9598 to 1.170 | 0.9933 to ??? |
| EC50 | 0.1614 to 0.2000 | 0.01272 to 0.01389 | 0.03238 to 0.04079 | 0.1298 to 0.1495 | 0.006321 to 0.008344 | 0.02935 to 0.03666 | 0.02130 to 0.07229 |
| Goodness of Fit | | | | | | | |
| Degrees of Freedom | 12 | 12 | 12 | 12 | 12 | 12 | 12 |
| R squared | 0.9950 | 0.9993 | 0.9961 | 0.9978 | 0.9935 | 0.9964 | 0.9054 |
| Sum of Squares | 97.79 | 17.82 | 87.63 | 47.63 | 141.7 | 78.75 | 3393 |
| Sy.x | 2.855 | 1.218 | 2.702 | 1.992 | 3.436 | 2.562 | 16.81 |
| Number of points | | | | | | | |
| # of X values | 14 | 14 | 14 | 14 | 14 | 14 | 14 |
| # Y values analyzed | 14 | 14 | 14 | 14 | 14 | 14 | 14 |

**Table S3. ELISA data for FGF2 binding to SDC1 ectodomains.**

| | $SDC1_{37}$ | $SDC1_{37HEP}$ | $SDC1_{37rHS}$ | $SDC1_{45,47}$ | $SDC1_{45,47HEP}$ | $SDC1_{45,47rHS}$ | $SDC1_{37CS}$ |
|---|---|---|---|---|---|---|---|
| log(agonist) vs. normalized response -- Variable slope | | | | | | | |
| Best-fit values | | | | | | | |
| LogEC50 | -1.570 | -2.007 | -1.936 | -1.535 | -1.773 | -1.789 | -1.789 |
| HillSlope | 1.307 | 1.073 | 0.8621 | 1.075 | 0.8626 | 0.8137 | 0.9714 |
| **$EC_{50}$** | 0.02690 | 0.009850 | 0.01158 | 0.02915 | 0.01687 | 0.01627 | 0.01627 |
| 95% CI (profile likelihood) | | | | | | | |
| LogEC50 | -1.596 to -1.545 | -2.105 to -1.906 | -2.076 to -1.793 | -1.596 to -1.474 | -1.890 to -1.654 | -1.950 to -1.624 | -1.875 to -1.702 |
| HillSlope | 1.225 to 1.398 | 0.8743 to 1.337 | 0.6784 to 1.114 | 0.9451 to 1.232 | 0.7079 to 1.065 | 0.6279 to 1.078 | 0.8270 to 1.153 |
| EC50 | 0.02537 to 0.02854 | 0.007848 to 0.01242 | 0.008387 to 0.01610 | 0.02534 to 0.03358 | 0.01287 to 0.02217 | 0.01122 to 0.02375 | 0.01334 to 0.01987 |
| Goodness of Fit | | | | | | | |
| Degrees of Freedom | 12 | 12 | 12 | 12 | 12 | 12 | 12 |
| R squared | 0.9988 | 0.9811 | 0.9669 | 0.9939 | 0.9781 | 0.9600 | 0.9875 |
| Sum of Squares | 27.33 | 330.8 | 535.0 | 127.2 | 377.7 | 671.5 | 229.3 |
| Sy.x | 1.509 | 5.250 | 6.677 | 3.256 | 5.610 | 7.481 | 4.371 |
| Number of points | | | | | | | |
| # of X values | 14 | 14 | 14 | 14 | 14 | 14 | 14 |
| # Y values analyzed | 14 | 14 | 14 | 14 | 14 | 14 | 14 |

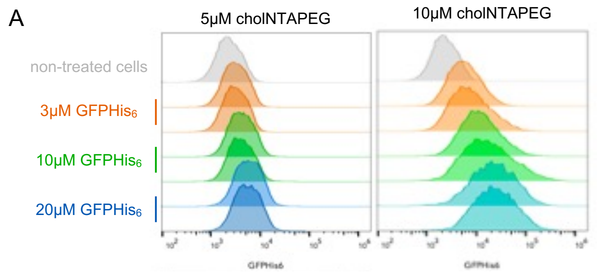


B

| sample | geometric mean (GFP) | average | MESF |
|---|---|---|---|
| 3 μM GFP | 16453 | | |
| 3 μM GFP | 16761 | **16607** | 512174.8 |
| 10 μM GFP | 54449 | | |
| 10 μM GFP | 46946 | **50697.5** | 649470.4 |
| 20 μM GFP | 40223 | | |
| 20 μM GFP | 63384 | **51803.5** | 653924.7 |

**Figure S18. Cell surface engineering with GFP-His$_6$ as a model.** (**A**)Flow cytometry analysis of live cells incubated with cholPEGNTA (5μM, left panel or 10μM, right panel; 1 hr, 37°C), followed by titrated incubations with GFP-His$_6$ (3, 10, or 20 μM; 1 hr, 37°C) to assess incorporation efficiency. Maximal fluorescence is observed at 10μM cholPEGNTA and 10μM GFP-His$_6$, demonstrating saturation of the cell surface with the His-tagged protein. (**B**) Using GFP calibration beads (TakaraBio # 632594) that report molecular equivalent soluble fluorochrome (MESF) units, we determined that for a 10 μM cholPEGNTA incubation, a subsequent incubation of GFPHis$_6$ at 3 μM is equivalent to ~512,000 MESF at the cell surface, 10 μM and 20 μM GFPHis$_6$ both result in ~650,000 MESF.

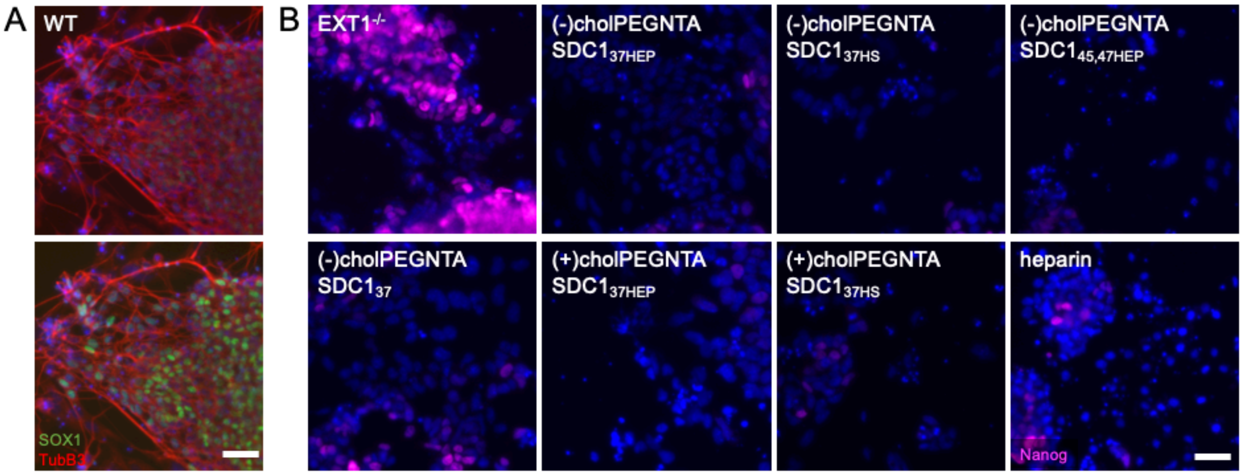


**Figure S19. Differentiation of adherent mESCs at D6. (A)** Representative images of WT mESCs before (top) and after (bottom) enhancement of SOX1 (green), as in **Fig 4D**. **(B)** Untreated EXT1$^{-/-}$ cells retain high Nanog expression, indicative of a pluripotent state. mESCs differentiated with SDC1 constructs or soluble heparin lose Nanog expression (pink) by D6. Scale bars: 50 μm.

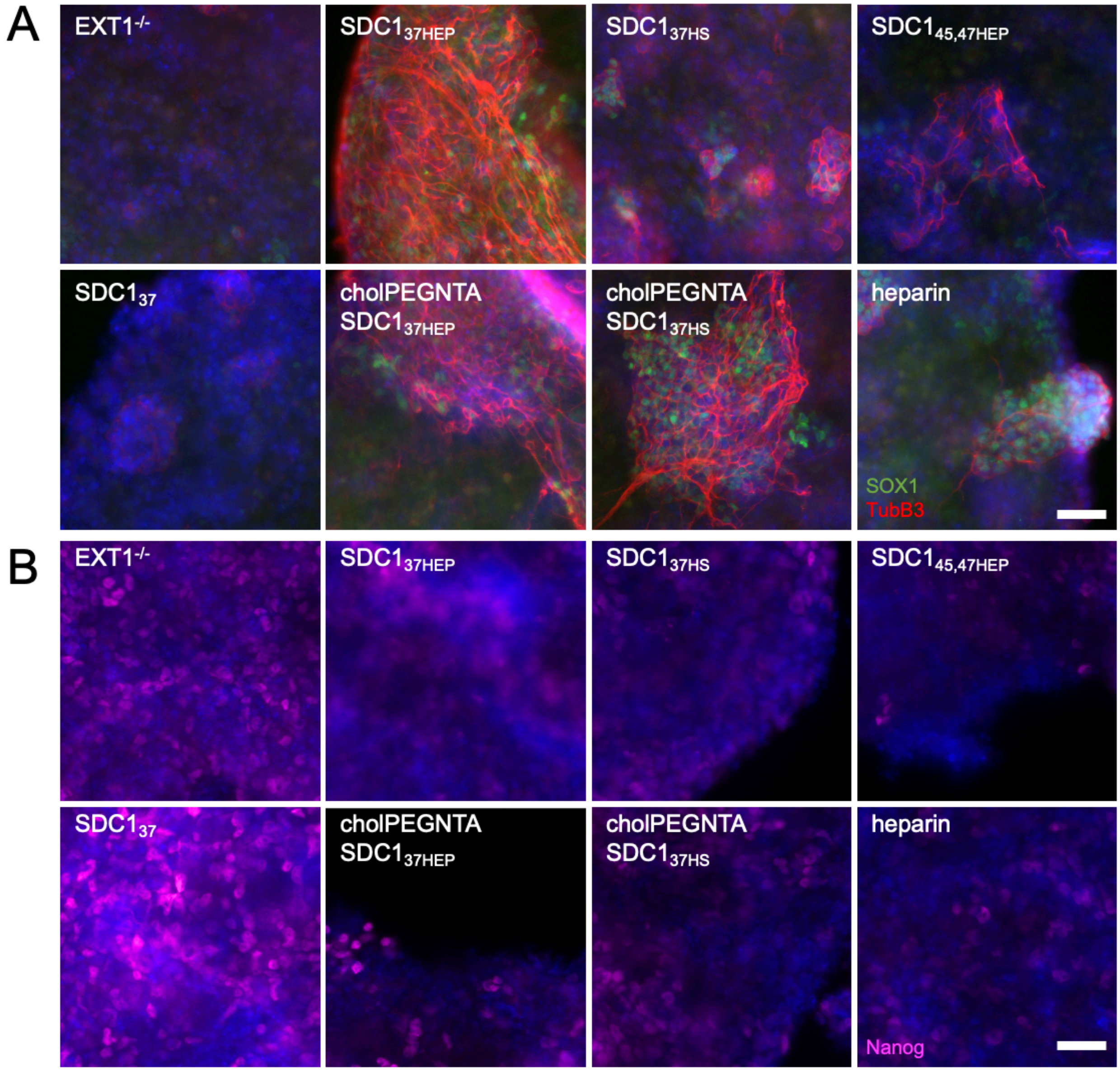


**Figure S20. Embryoid body (EB) differentiation at D6. (A)** After 6 days of differentiation, EXT1$^{-/-}$ EBs remodeled with GAG-conjugated SDC1 constructs, or soluble heparin, differentiated into neuronal precursor cells as indicated by SOX1$^{+}$ (green) cells and βIII tubulin (TubB3, red) expression. **(B)** Concurrent with increased differentiation markers, Nanog (pink) is decreased in cells treated with soluble heparin or replete SDC1 constructs. Untreated cells and those treated with SDC1$_{37}$ core protein alone retain high Nanog expression. Scale bars: 50 μm.

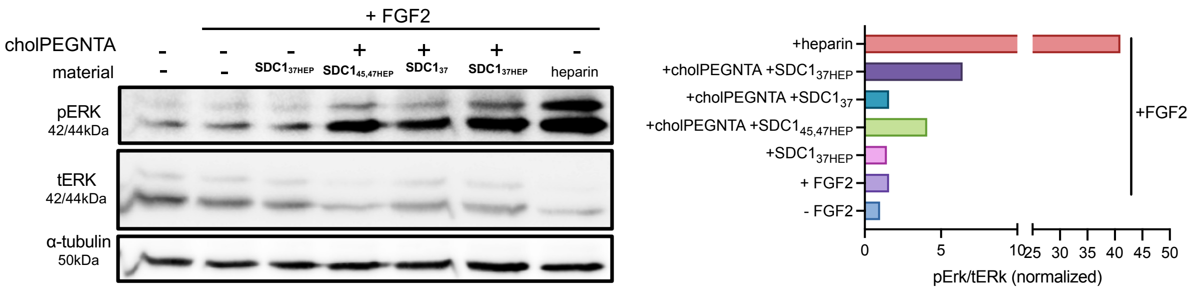


**Figure S21. FGF2 triggers the activation of the ERK pathway.** When stimulated with FGF2 (15 mins, 37°C), the addition of soluble heparin or membrane bound SDC1 constructs resulted in increased downstream ERK phosphorylation. Ext1$^{-/-}$ mESCs were pre-incubated with 2 μM of SDC1 constructs, with or without prior treatment with cholPEGNTA. Excess was washed away before incubation with FGF2 in full media. This treatment is in stark contrast with heparin treated cells, in which soluble heparin was added to the FGF2-supplemented media at D0-D2.

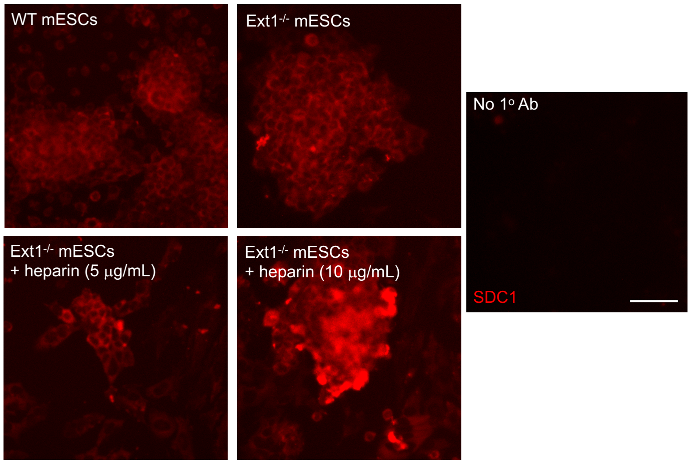


**Figure S22. mESCs express SDC1.** Cells were stained for SDC1 using anti-SDC1 antibody (clone DL-101) after 6 days of differentiation in N2B27 media.

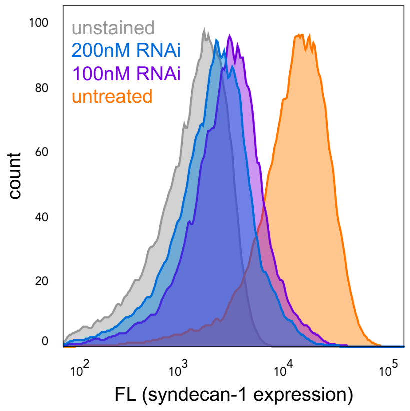


**Figure S23.** MDA-MB-231 cells treated with 100 (purple) or 200nM (blue) pooled SDC1 dsRNAi exhibit reduced SDC1 expression compared to non-targeting DsiRNA control (orange). Unstained cells (gray) are those incubated with secondary antibody only.

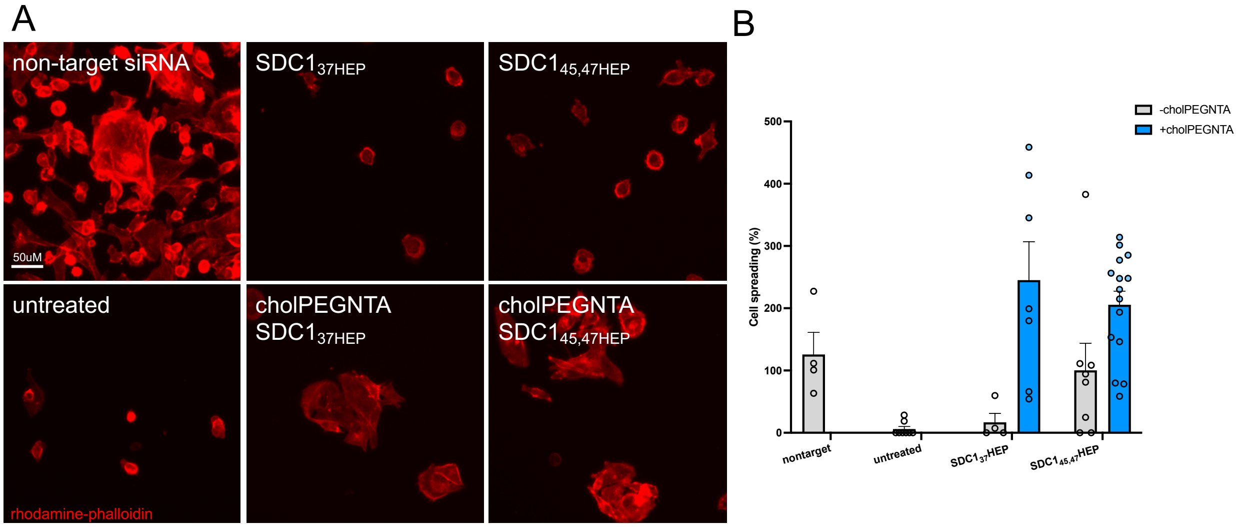


**Figure S24. Rescue of cell spreading in SDC1$^{KO}$ cells.** (**A**) Immunofluorescence imaging of phalloidin-stained SDC1$^{KO}$ MDA-MB-231 cells following cell spreading (2 hr) on vitronectin matrices show that SDC1$_{37HEP}$ and SDC1$_{45,47HEP}$ rescue cell spreading only when they are anchored onto cell surfaces. (**B**) Quantification of cell spreading.

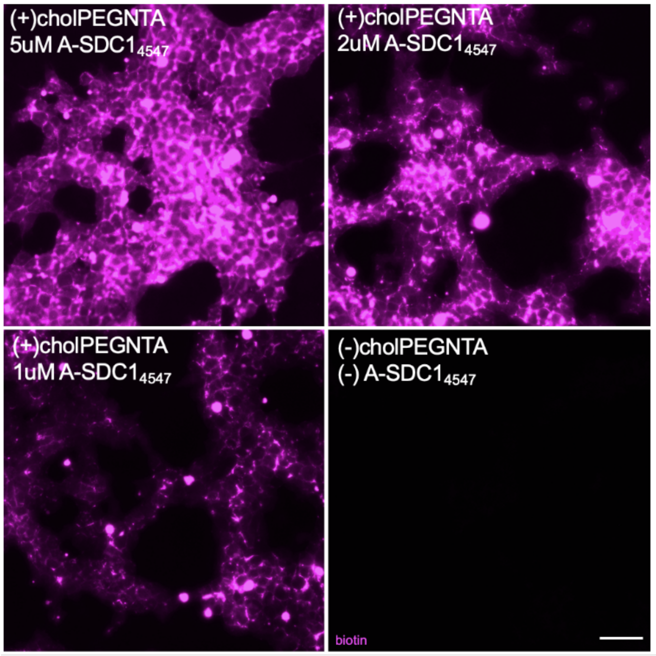


**Figure S25. Proximity tagging across different concentrations.** Live WT mESCs pre-treated with cholPEGNTA (10 μM, 1 hr) were incubated with A-SDC1$_{45,47}$ at various concentrations (5, 2, 1 μM) for 1 hr, 37$^{o}$C. Proximity tagging was then initiated to biotinylate the interactors of A-SDC1$_{45,47}$. The interactors were then probed using a fluorophore-conjugated streptavidin (pink).

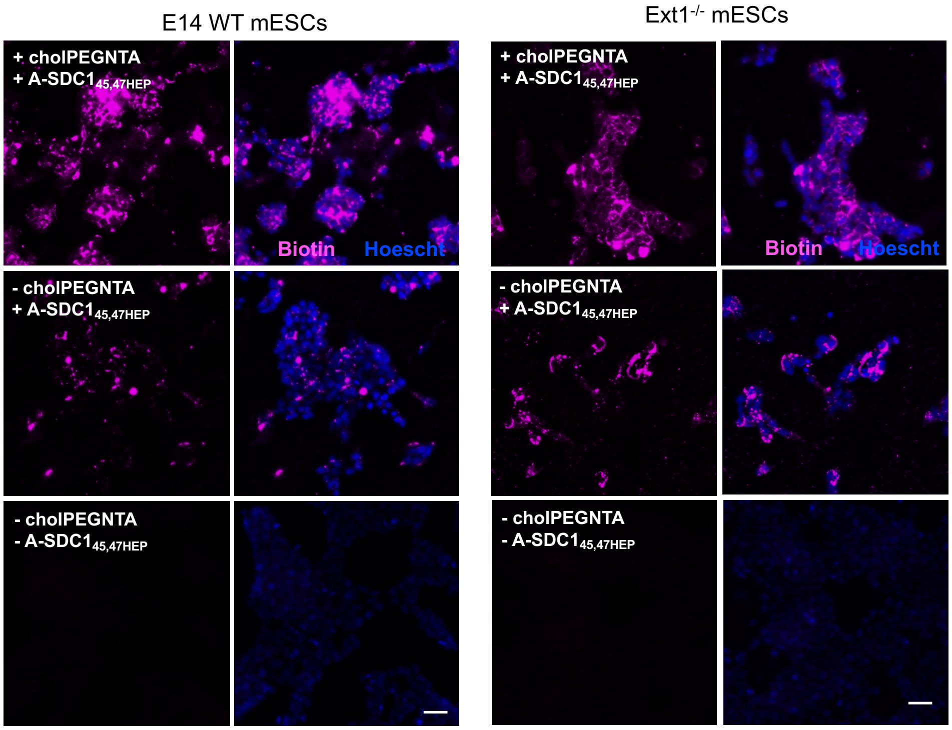


**Figure S26. Proximity tagging in WT and Ext1$^{-/-}$ mESCs.** Biotinylated interactors are detected by Cy5-streptavidin (pink). Treatment with soluble A-SDC1$_{45,47HEP}$ resulted in less fluorescence detection than membrane-bound in both WT and EXT1$^{-/-}$ mESCs.
